# Venom vesicles from the parasitoid *Ganaspis hookeri* facilitate venom protein entry into host immune cells

**DOI:** 10.1101/2025.05.08.652939

**Authors:** Nicholas M. Bretz, Chris Lark, Sarah Perkel, Rebecca Jackson, Jeremy D. Driskell, James A. Mobley, Nathan T. Mortimer

## Abstract

The parasitoid wasp *Ganaspis hookeri* infects *Drosophila melanogaster* larvae, laying an egg and injecting venom directly into the host body cavity. While infected hosts mount an immune response in an attempt to eliminate the parasitoid egg, parasitoid venom proteins act to inhibit these host immune responses and manipulate host physiology to ensure infection success. A key immune suppressive venom protein in *G. hookeri* is a venom- specific isoform of the SERCA (<u>S</u>arco/<u>e</u>ndoplasmic <u>r</u>eticulum <u>C</u>a^2+^-<u>A</u>TPase) calcium pump. However, SERCA is a large hydrophobic protein, and the mechanism by which it and other venom proteins are transported into the host is not well understood. We used a variety of biophysical, biochemical, and cell biological approaches to assess the properties of *G. hookeri* venom. Electronic microscopy and nanoparticle tracking analysis revealed the presence of venom vesicles as a putative transport mechanism. We used tunable resistive pulse sensing (TRPS) to biophysically characterize these vesicles, and our TRPS data suggest *G. hookeri* venom is composed of multiple vesicle types that are distinguishable by size, zeta potential, and density. Finally, we fluorescently labeled venom vesicles to test for entry into host immune cells. We observed that these vesicles interact with immune cells membranes and are internalized into the cell. Our data support a model in which *G. hookeri* venom proteins, including SERCA, are packaged into an array of venom vesicles, and transported into host cells for immune suppression.

## Introduction

Parasitoid wasps comprise an ecologically diverse group of insects numbering an estimated 350,000 species worldwide (Saunders and Ward, 2018). These parasitoids infect a wide range of insect host species, and while virulence strategies differ dependent on the host and parasitoid species, wasps use an array of virulence factors, including venom proteins, calyx proteins, and symbiont polydnaviruses to overcome host defense responses and ensure offspring fitness (Asgari et al., 1998; Beckage, 1998; Moreau and Asgari, 2015). These parasitoid virulence factors target host immunity using a wide range of mechanisms including the destruction of immune cells or immune cell precursors (Ramroop et al., 2021), dysregulation of the immune cell actin cytoskeletal structure (Labrosse et al., 2005), as well as altering the adhesion of immune cells important in the encapsulation process (Strand, 1994). Parasitoid venoms also enhance offspring fitness through manipulation of host development and metabolism, including altering expression of host genes associated with glycolysis and gluconeogenesis (Martinson et al., 2014), and changing activity of host metabolic enzymes and pathways (Song et al., 2022; Kryukova et al., 2024).

The parasitoid *Ganaspis hookeri* is a generalist of *Drosophila* species, with high infection success against *Drosophila melanogaster* (Mortimer et al., 2013). *G. hookeri* is an endoparasitoid of *Drosophila* larvae, and during infection, lays an egg and injects virulence factor containing venom into the host hemocoel. Following parasitoid infection, *D. melanogaster* larvae mount a conserved cellular immune response against the wasp egg (Mortimer, 2013; Kim-Jo et al., 2019). This response is initiated by a ‘calcium burst’, leading to the calcium-signaling mediated activation of plasmatocytes (Mortimer et al., 2013), a class of circulating hemocytes with immune functions similar to mammalian macrophages (Hultmark and Andó, 2022). Activation enables plasmatocytes to migrate towards and bind to the wasp egg, forming an inner capsule that is later encapsulated by lamellocytes, a parasitoid induced hemocyte class, and melanized leading to parasitoid death (Russo et al., 1996; Nappi and Christensen, 2005; Mortimer et al., 2012).

*G. hookeri* venom consists of a complex mix of proteins with a wide range of predicted functions (Mortimer et al., 2013; Alvarado et al., 2020). We have shown that *G. hookeri* infection inhibits host plasmatocyte activation utilizing a venom-specific isoform of the Sarco-endoplasmic reticulum calcium ATPase (SERCA) (Mortimer et al., 2013). The SERCA protein is widely conserved, and acts as an active calcium pump, removing Ca2+ ions from the cytoplasm into SR/ER stores (Altshuler et al., 2012). Our previous work showed that *G. hookeri* infection antagonizes the plasmatocyte calcium burst, and that this activity is dependent on venom SERCA activity, suggesting that venom SERCA interacts with host cells to lower cytoplasmic calcium levels (Mortimer et al., 2013).

Venom SERCA is a 110 kDa protein with 11 predicted transmembrane domains, which leads to an interesting problem: our results suggest that venom SERCA, despite being a large, hydrophobic protein, is transported through aqueous environments of the venom and host hemocoel before it enters host cells to manipulate calcium signaling. Other parasitoid species with a similar venom protein composition to *G. hookeri*, including large hydrophobic transmembrane proteins, appear to have solved this problem through the evolution of venom extracellular vesicles (EVs) (Gueguen et al., 2011; Goecks et al., 2013; Heavner et al., 2013, 2017; Wan et al., 2019; Ramroop et al., 2021).

*Leptopilina heterotoma*, a generalist parasitoid of *Drosophila* species, utilizes EVs, formerly referred to as virus-like particles, to infect *Drosophila* hosts. *L. heterotoma* EVs are largely homogenous, with an estimated size of 300 nm and displaying a knob-like morphology (Morales et al., 2005; Chiu et al., 2006; Heavner et al., 2017). These *L. heterotoma* EVs interact with host hemocytes to induce lamellocyte lysis and cell death within the hematopoietic lymph gland (Rizki and Rizki, 1990; Ramroop et al., 2021). Similarly, the venom of *Leptopilina boulardi*, a specialist parasitoid of melanogaster subgroup larvae, also contains extracellular vesicles more commonly known as venosomes (Gueguen et al., 2011; Wan et al., 2019). These vesicles are less well characterized but exhibit sizes ranging from 100 nm - 200 nm as measured through imaging by electron microscopy (Wan et al., 2019). These vesicles also appear aggregation-prone, and are often imaged in aggregates of 400 nm - 1 µm (Wan et al., 2019). *L. boulardi* employs a RhoGAP protein that is predicted to disrupt actin cytoskeletal architecture and adhesion properties of host hemocytes, and this protein is found within *L. boulardi* venosomes (Colinet et al., 2007; Wan et al., 2019). We therefore hypothesize that *G. hookeri* venom contains vesicles to enable the transport of venom SERCA and associated venom proteins during infection. To test our hypothesis, we utilized a combination of biochemical, biophysical, and cell biological assays to characterize the contents of *G. hookeri* venom.

## Results

### SERCA is expressed in *G. hookeri* venom gland cells

To establish the localization of SERCA expression in *G. hookeri* venom producing cells, whole venom apparatuses from female *G. hookeri* wasps were dissected, stained with anti-SERCA and labeled phalloidin, and imaged using confocal microscopy (Figure 1A-C; bright field Figure S1). Maximum intensity projections of Z-stack images taken at 40x magnification reveal that SERCA is expressed in cells of the venom gland (Figure 1A; indicated by the arrow), and is also found within the venom reservoir (Figure 1A; indicated by the arrowhead). SERCA localization appears to be non-random within the venom gland, but rather shows a distinct localization pattern throughout the gland. To better characterize this SERCA localization, stained venom glands were imaged at 63x magnification. Z-stack images were projected through three dimensions, and 2D slices of the projection were analyzed for SERCA localization (Figure 1D-F). The projection shows that the SERCA stain appears to associate with actin structures within the venom gland (Figure 1D-F; indicated by arrow). The *L. heterotoma* venom gland is organized around actin channels serving to transport venom proteins for secretion (Ferrarese et al., 2009). This organization allows *L. heterotoma* venom proteins to be packaged into extracellular vesicles, and so we hypothesized that SERCA might be similarly packaged in *G. hookeri* venom glands.

**Figure 1.**
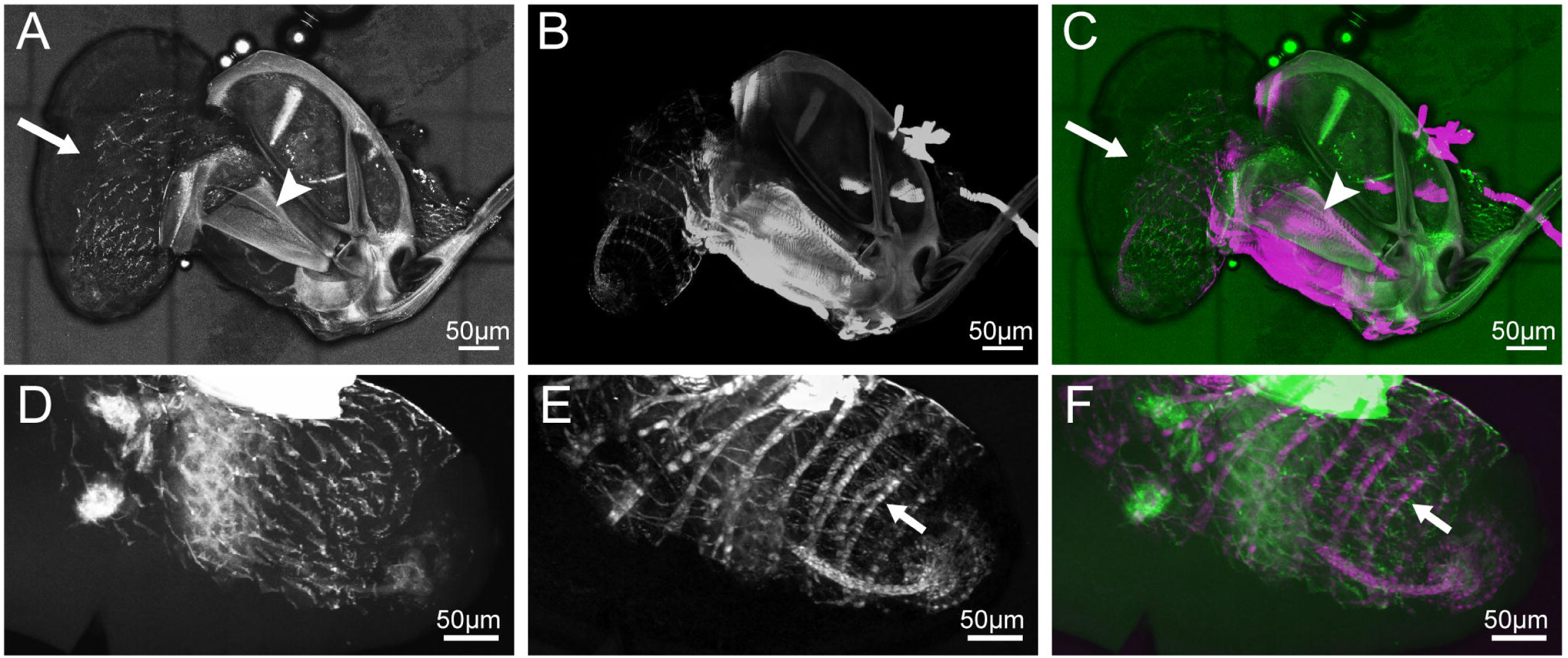
Confocal micrographs of SERCA staining in *G. hookeri* venom apparatuses. (A-C) Maximum projection of Z-stack images taken at 40x magnification. (A) anti-SERCA, (B) phalloidin-Alexa Fluor 633, and (C) merged. Arrows in A, C indicate the venom gland and arrowheads in A, C indicate the venom reservoir. (D-F) 3D projection of Z-stack images taken at 63x magnification. (D) anti-SERCA, (E) phalloidin-Alexa Fluor 633, and (F) merged. Arrows in E-F indicate actin structures. Scale bars are 50 µm in each panel.

### Characterization of *G. hookeri* venom reveals a heterogenous population of venom vesicles

To determine whether *G. hookeri* venom contains vesicles, purified whole venom was characterized with scanning electron microscopy (SEM; Figure 2A) and negative stain transmission electron microscopy (TEM; Figure 2B). We observed a variety of particles in venom, with different sizes and morphologies. Many of the particles are <40 nm, and are unlikely to be membrane enclosed vesicles (Figure 2A, example indicated by the arrow). We also observe a range of particles with sizes between 40-200 nm that appear to correspond to vesicles (Figure 2A, examples indicated by arrowheads). Based on size, these venom vesicles can best be characterized as small extracellular vesicles (<200 nm; sEVs) (Théry et al., 2018). These venom sEVs are considerably smaller than the venom vesicles described in *L. heterotoma*, and polydisperse, rather than the more uniform distribution of venom vesicles observed in both *L. heterotoma* and *L. boulardi*. These observations further support the conclusion that *G. hookeri* venom contains vesicles, and suggests that *G. hookeri* venom contains a heterogenous mixture of vesicles.

**Figure 2.**
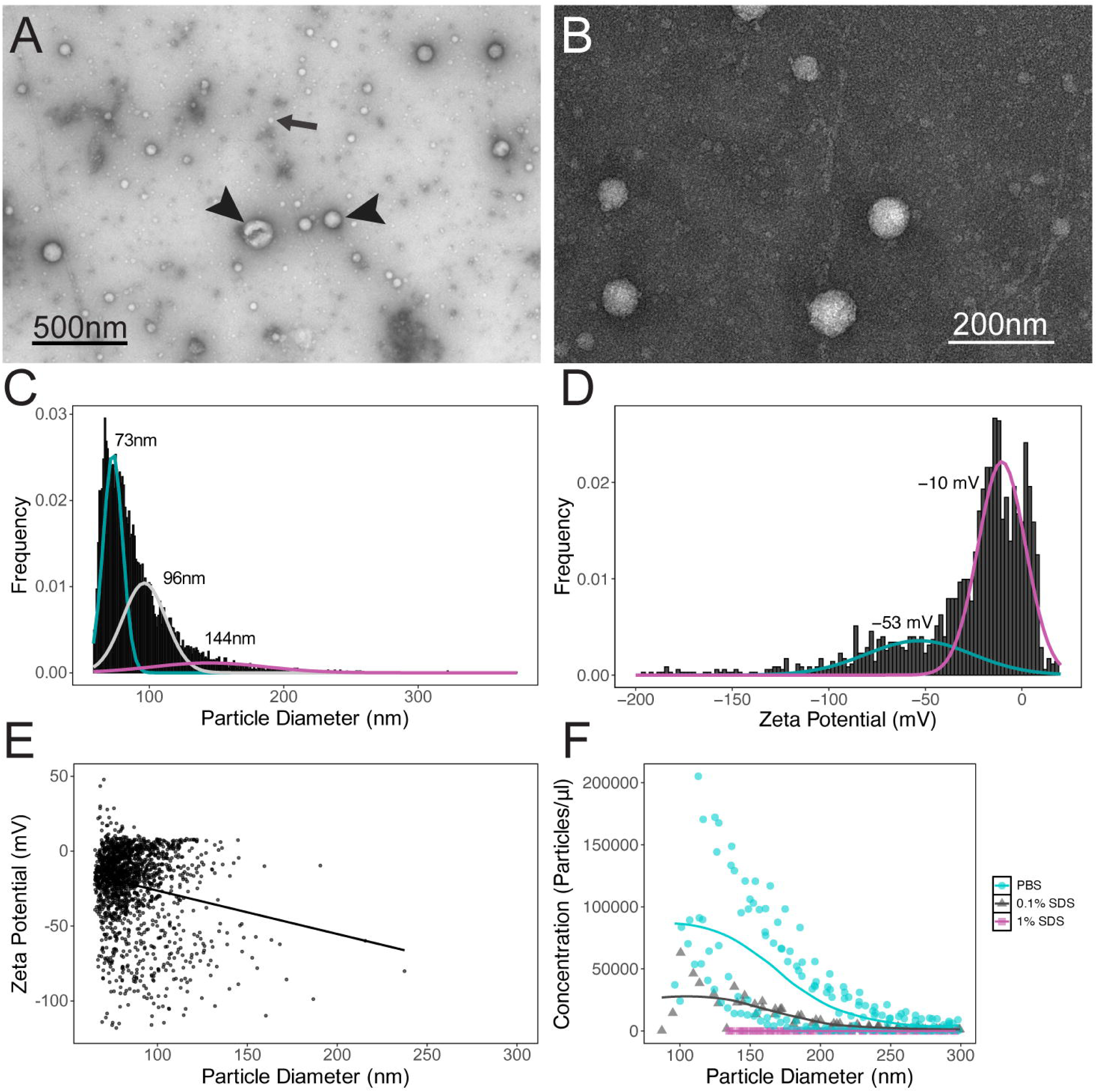
Characterization of vesicles in whole venom. (A) Scanning electron micrograph of whole venom. Arrow indicates a small non-vesicle particle, arrowheads indicate differently sized vesicles. Scale bar is 500 nm. (B) Transmission electron micrograph of whole venom. Scale bar is 200 nm. (C) Histogram of vesicle diameter measured by TRPS. Mixture model distributions are overlaid, with each distribution mean indicated. (D) Histogram of vesicle zeta potential measured by TRPS. Mixture model distributions are overlaid, with each distribution mean indicated. (E) Scatter plot of vesicle diameter vs zeta potential with calculated linear regression line. (F) Scatter plot of vesicle diameter vs concentration following each of the indicated detergent treatments, with calculated loess local weighted regression lines.

To characterize *G. hookeri* venom vesicles, we used tunable resistive pulse sensing (TRPS) to measure venom vesicle size and concentration. The distribution of the corresponding histogram of measured particle diameters (Figure 2C) further supports our conclusion that *G. hookeri* venom vesicles display polydispersity. By using mixture modeling to estimate subpopulations within a putatively multimodal distribution, we uncovered the likelihood of three distinct vesicle subpopulations by size, corresponding to a subpopulation with a mean (µ) of 72.9 ± the standard deviation (σ) of 7.2 nm and accounting for 46.1% of measured vesicles (weight or proportion of observations; λ), a second subpopulation with a mean of 96.2 ± 16.0 nm and accounting for 41.9% of measured vesicles, and a third subpopulation with a mean of 144.5 ± 41.9 nm and accounting for 12.1% of measured vesicles.

TRPS was next used to measure the zeta potential (or surface charge) of venom vesicles. We observed a similar wide range of measured zeta potentials among vesicles (Figure 2D). Mixture modeling supports the existence of two subpopulations, the first of which has a mean zeta potential of -10.4 ± 12.3 mV and accounting for 68.2% of measured vesicles, and a second subpopulation with a mean zeta potential of -53.2 ± 27.5 mV and accounting for 24.6% of measured vesicles (with 7.2% of vesicles unaccounted for by these modeled distributions). To explore the relationship between vesicle diameter and zeta potential, we plotted vesicle size vs zeta potential and calculated the linear regression line (Figure 2E). Correlation analysis suggests a weak, yet significant, negative correlation between size and zeta potential (Pearson’s r = -0.19, p = 6.4e-15), suggesting that larger vesicles may have increasingly negative surface charge. However, the scatter plot (Figure 2E) and small Pearson’s r suggests that the relationship between vesicle size and zeta potential is likely more complex.

To characterize the composition of the vesicles, we tested their sensitivity to detergent. We incubated whole venom with increasing concentrations of sodium dodecyl sulfate (SDS), a strong ionic detergent that would be predicted to disrupt vesicle lipid organization, and analyzed vesicle size and concentration by TRPS. PBS control treated venom shows the expected histogram distribution (Figure 2F, cyan). Incubation with 0.1% SDS led to a noticeable decrease in vesicles of all sizes (Figure 2F, charcoal). Finally, incubation with 1% SDS abolished nearly all vesicles found in *G. hookeri* venom (Figure 2F, magenta).

### Density gradient ultracentrifugation resolves *G. hookeri* venom into distinct fractions

To resolve the complex composition of *G. hookeri* venom, we used discontinuous sucrose density gradient fractionation of whole venom. To test the effectiveness of gradient fractionation for producing unique subpopulations of venom proteins, we performed sodium dodecyl sulfate-polyacrylamide gel electrophoresis (SDS-PAGE) analysis to determine the protein composition of whole venom compared to each fraction (Figure 3A). Whole venom (Figure 3A, WV) shows a complex banding pattern, which is simplified within each fraction. The venom fractions also display discrete banding patterns, suggesting that their protein contents are distinct from each other (Figure 3A, F5-F1). The least dense fractions (Figure 3A, F2-F1) appear to have a much lower protein content, suggesting that most proteins are associated with higher density complexes.

**Figure 3.**
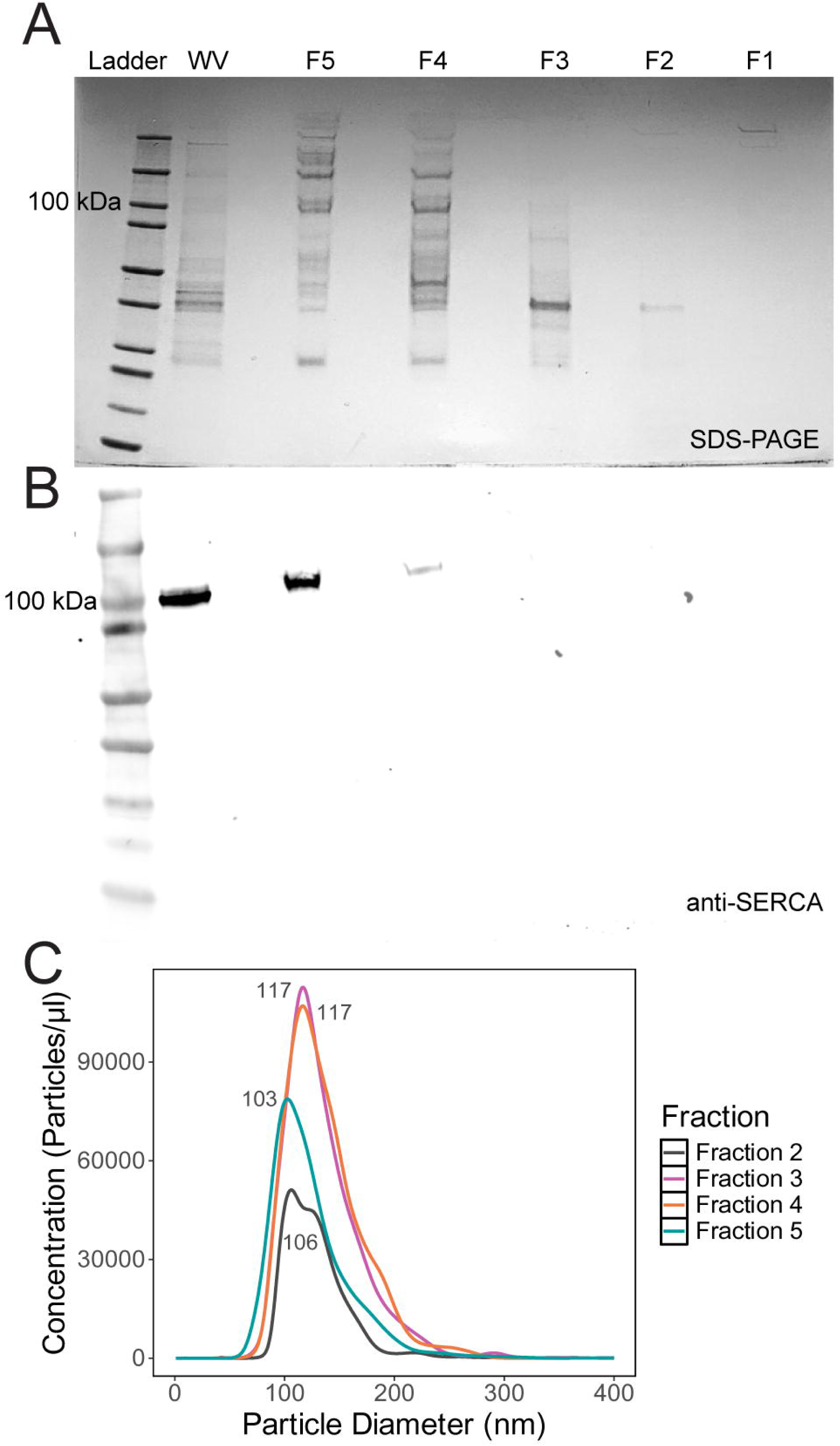
Sucrose gradient fraction of *G. hookeri* venom. (A) SDS-PAGE analysis of whole venom (WV) and each indicated fraction. (B) Anti-SERCA Western blot analysis of whole venom following SDS-PAGE as shown in (A). (C) Distribution of vesicle diameter by concentration within each indicated fraction as measured by nanoparticle tracking analysis.

Because we hypothesize that SERCA is likely found in venom vesicles, we performed Western blot analysis to determine which venom fraction contains SERCA. The Western blot shows SERCA in whole venom as expected (Figure 3B, WV). We also observe that SERCA is predominantly found in the most dense fraction (F5), with a small amount of protein also visible in F4. This suggests that vesicles are likely found predominantly in F5, which would be unexpected given that F5 represents the 50% sucrose fraction with a density of 1.23 g/mL (Hofmann, 1977). Typical extracellular vesicles, including exosomes and microvesicles, have densities in the range of 1.08-1.19 g/mL (Brennan et al., 2020), which would better correspond to fractions F2-F4 (1.08-1.18 g/mL) (Hofmann, 1977) in our gradient. To test whether vesicles were also found in these fractions, we used nanoparticle tracking analysis (NTA) to analyze vesicles across all fractions (Figure 3C). We could not detect vesicles in F1, but vesicles were observed in all other fractions. Interestingly, F2 and F5 had a similar predominant size peak although the overall curves were differently shaped. F2 and F5 were separable from F3 and F4 which showed identical peak sizes and remarkably similar curves. Based on these observations, we hypothesize that venom vesicles are found at distinct densities, and that these between-fraction differences are not due solely to particle size, given the similarities between vesicles found in F2 and F5, which represent dissimilar densities (1.08 g/mL and 1.23 g/mL; Figure 3C).

### Biophysical characterization of venom vesicles suggests discrete vesicle populations

To better understand the relationship between fractions, we performed in-depth imaging and biophysical characterization. Based on our NTA data (Figure 3C), we first focused on the relationship between F2 and F5. Negative stain TEM revealed that the vesicle contents of F2 are largely monodisperse in size and display a consistent round morphology (Figure 4A). Morphologically similar vesicles are also observed in F5 (Figure 4B), however, F5 also contains a class of larger vesicles with a distinct “kidney bean” shaped morphology (Figure 4C). Negative stain TEM for F5 characterization was performed on unfixed samples to avoid possible morphology artefacts associated with fixation (Kellenberger et al., 1992).

**Figure 4.**
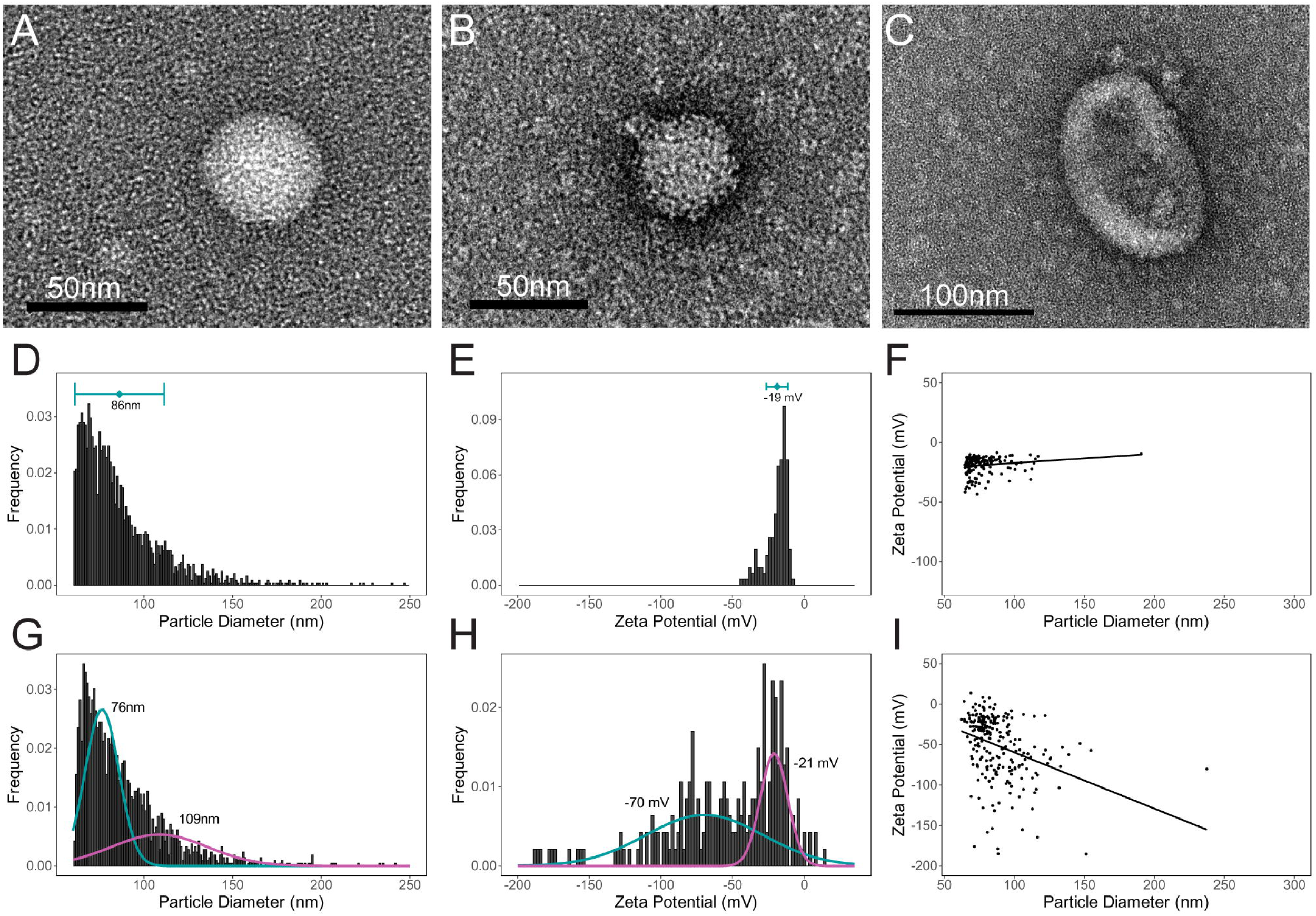
Biophysical characterization of fractions F2 and F5. (A-C) Transmission electron micrographs of vesicles from F2 (A) and F5 (B-C). Scale bars are 50 nm (A-B) and 100 nm (C). (D) Histogram of F2 vesicle diameter measured by TRPS with the mean ± standard deviation of the vesicle population indicated. (E) Histogram of F2 vesicle zeta potential measured by TRPS with the mean ± standard deviation of the vesicle population indicated. (F) Scatter plot of F2 vesicle diameter vs zeta potential with calculated linear regression line. (G) Histogram of F5 vesicle diameter measured by TRPS. Mixture model distributions are overlaid, with each distribution mean indicated. (H) Histogram of F5 vesicle zeta potential measured by TRPS. Mixture model distributions are overlaid, with each distribution mean indicated. (I) Scatter plot of F5 vesicle diameter vs zeta potential with calculated linear regression line.

We performed TRPS in order to more thoroughly characterize the composition of each fraction. In agreement with the TEM imaging results, F2 has a relatively monodisperse population of vesicles (Figure 4D-F). Vesicles in F2 have a mean diameter of 86.2 ± 25.2 nm (Figure 4D), and mean zeta potential of -19.0 ± 7.4 mV (Figure 4E). The linear regression line of the size vs zeta potential plot (Figure 4F) shows a weak positive correlation between these properties (r = 0.16, p = 9.3e-5).

Conversely, TPRS analysis of the vesicles in F5 presents evidence for polydispersity, in agreement with the TEM imaging results. Due to this evidence of a mixed population, we performed mixture modeling of vesicle size data. Our modeling suggests that F5 contains two populations of vesicles (Figure 4G); the first subpopulation has a mean diameter of 76.4 ± 9.5 nm and accounts for 63.6% of the vesicles measured, and the second subpopulation has a mean diameter of 108.7 ± 27.2 nm and accounts for 36.3% of the vesicles measured. This bimodal distribution is also reflected in our measurements of vesicle zeta potential, with the mixture model again suggesting two populations (Figure 4H). The first subpopulation has a mean zeta potential of -69.7 ± 40.9 mV and accounts for 66.0% of the vesicles measured, and the second subpopulation has a mean zeta potential of -21.1 ± 9.5 mV and accounts for 34.0% of the vesicles measured. Linear regression of the size vs charge plot (Figure 4I) suggests that there is a moderate negative correlation (r = -0.34, p = 2.2e-16) with the larger population of vesicles having a more negative zeta potential.

Our NTA data (Figure 3C) further suggest that F3 and F4 likely contain highly similar vesicle populations. Negative stain TEM imaging supports this hypothesis with both fractions showing largely homogenous populations of similarly sized, round vesicles (Figure 5A-B). TRPS analysis bears out the morphological similarity, with F3 having a mean diameter of 96.5 ± 32.9 nm and F4 having a mean diameter of 93.4 ± 31.6 nm (Figure 5C-D). However, characterization of zeta potential suggests that F3 and F4 are distinct despite this morphological similarity (Figure 5E-F). F3 has a mean zeta potential of -12.5 ± 26.9 mV. Despite the homogeneity of size and morphology, F4 appears to have a bimodal distribution of zeta potentials; the first subpopulation has a mean zeta potential of -65.1 ± 47.3 mV and accounts for 51.1% of the vesicles measured, and the second subpopulation has a mean zeta potential of -22.4 ± 10.6 mV and accounts for 48.9% of the vesicles measured (Figure 5F). Linear regression of the size vs charge plots suggests that there is no correlation (r = -0.031, p = 0.062) between size and zeta in F3 (Figure 5G), and a weak correlation (r = -0.3, p = 2.2e-16) in F4 (Figure 5G).

**Figure 5.**
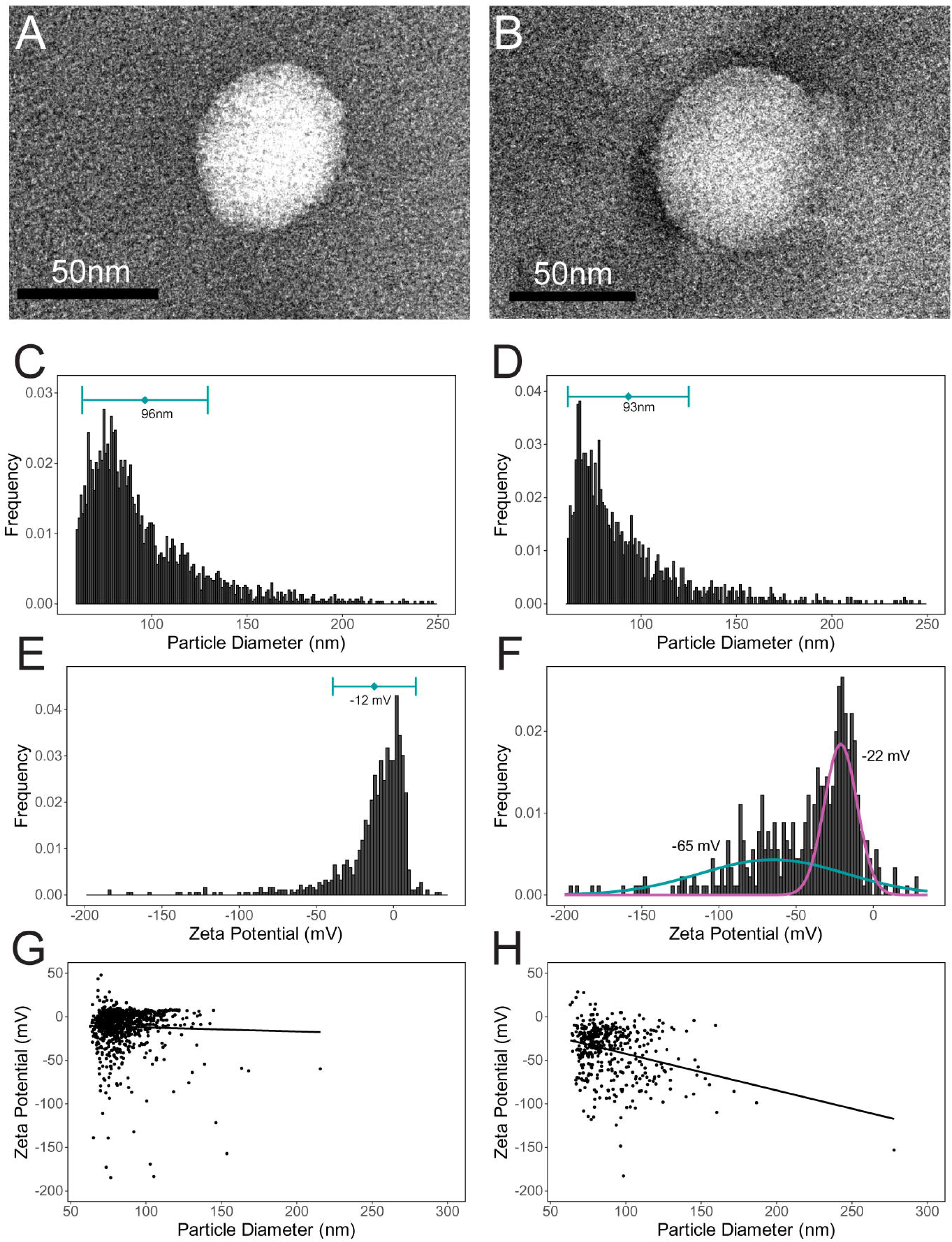
Biophysical characterization of fractions F3 and F4. (A-B) Transmission electron micrographs of vesicles from F3 (A) and F4 (B). Scale bars are 50 nm. (C) Histogram of F3 vesicle diameter measured by TRPS with the mean ± standard deviation of the vesicle population indicated. (D) Histogram of F4 vesicle diameter measured by TRPS with the mean ± standard deviation of the vesicle population indicated. (E) Histogram of F3 vesicle zeta potential measured by TRPS with the mean ± standard deviation of the vesicle population indicated. (F) Histogram of F4 vesicle zeta potential measured by TRPS. Mixture model distributions are overlaid, with each distribution mean indicated. (G) Scatter plot of F3 vesicle diameter vs zeta potential with calculated linear regression line. (H) Scatter plot of F4 vesicle diameter vs zeta potential with calculated linear regression line.

### Mass spectrometry identifies the protein composition of *G. hookeri* venom fractions

Our observation of venom vesicles in *G. hookeri* venom suggests that they serve to transport venom proteins into host cells following infection as has been observed in other parasitoid species. However, in these other species, venom proteins are found within a single vesicle type (Heavner et al., 2017; Wan et al., 2019), rather than the heterogeneous population of venom vesicles we have characterized in *G. hookeri*. We hypothesize that the distinct vesicle populations we observe across venom fractions may contain distinct protein cargos. To test this, we performed mass spectrometry on proteins purified from each fraction, and mapped the resulting peptides onto the *G. hookeri* transcriptome to identify the proteins found in each venom fraction.

After filtering our results to determine fraction membership, we identified 98 proteins in F1, 128 proteins in F2, 118 in F3, 114 in F4, and 94 in F5. We observe considerable overlap between fractions (Figure 6A). However, each fraction does have unique members (selected examples shown in Table 1), suggesting that there is a potential basis for vesicle or fraction specific functions. Among these unique proteins is SERCA, which was specifically found in F5, in accordance with our Western blot data (Figure 3B). We also observe a trend that the molecular weight and isoelectric point of venom proteins increase with increasing density, although a wide range of values for each of these protein characteristics are found in each fraction (Figure 6B-C). The molecular weight data determined from our mass spectrometry results (Figure 6B) is generally in agreement with our fraction specific SDS-PAGE protein banding patterns (Figure 3A), which also illustrate a trend toward larger proteins in F4 and F5.

**Figure 6.**
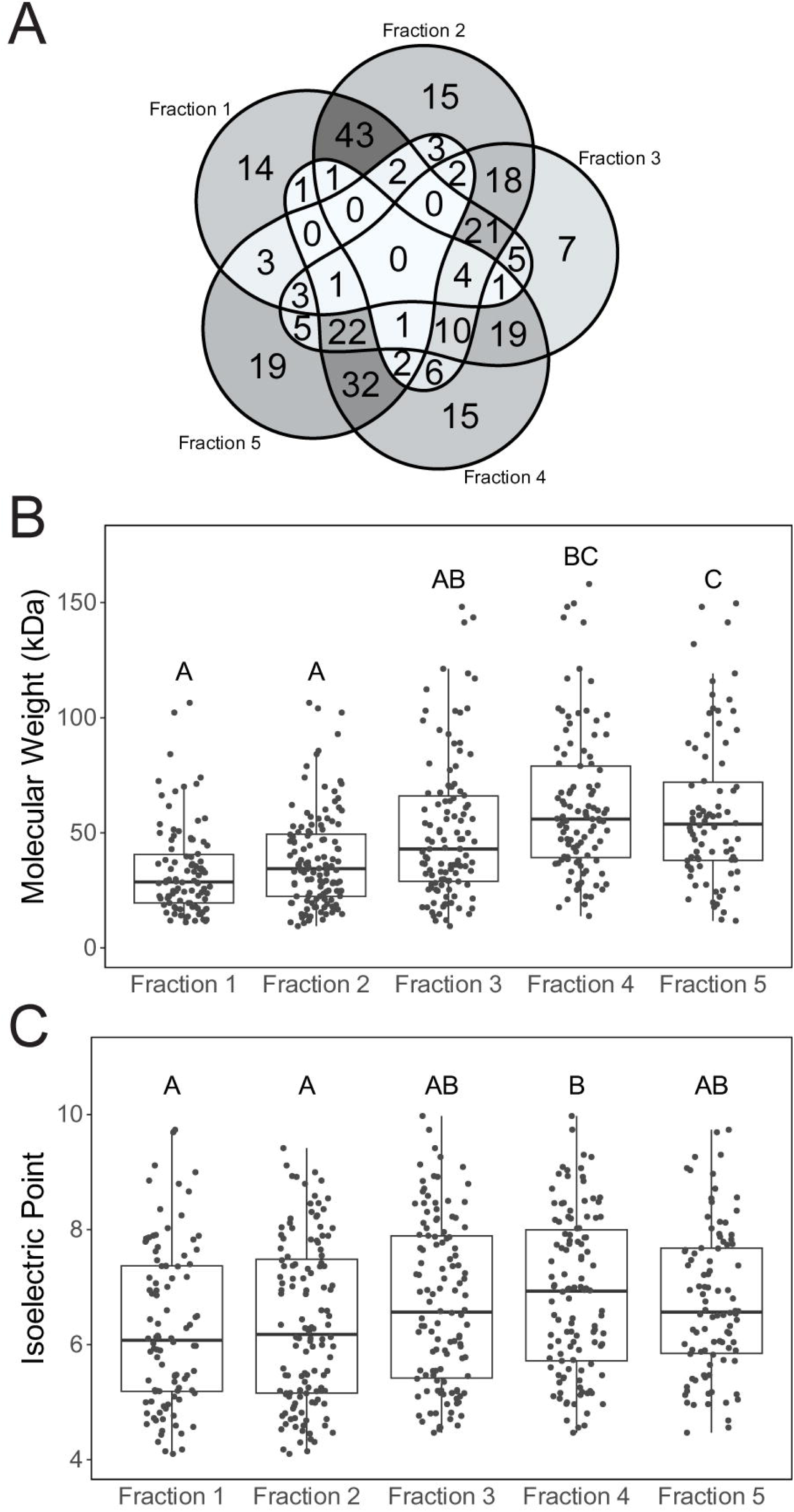
Fraction-specific mass spectrometry. (A) Venn diagram of fraction membership of all proteins identified by mass spectrometry. (B) Box plot of protein molecular weight by fraction with individual data points overlaid. Letters indicate confidence groups assigned by Tukey’s HSD *post hoc* testing. (C) Box plot of protein isoelectric point by fraction with individual data points overlaid. Letters indicate confidence groups assigned by Tukey’s HSD *post hoc* testing.

**Table 1.**
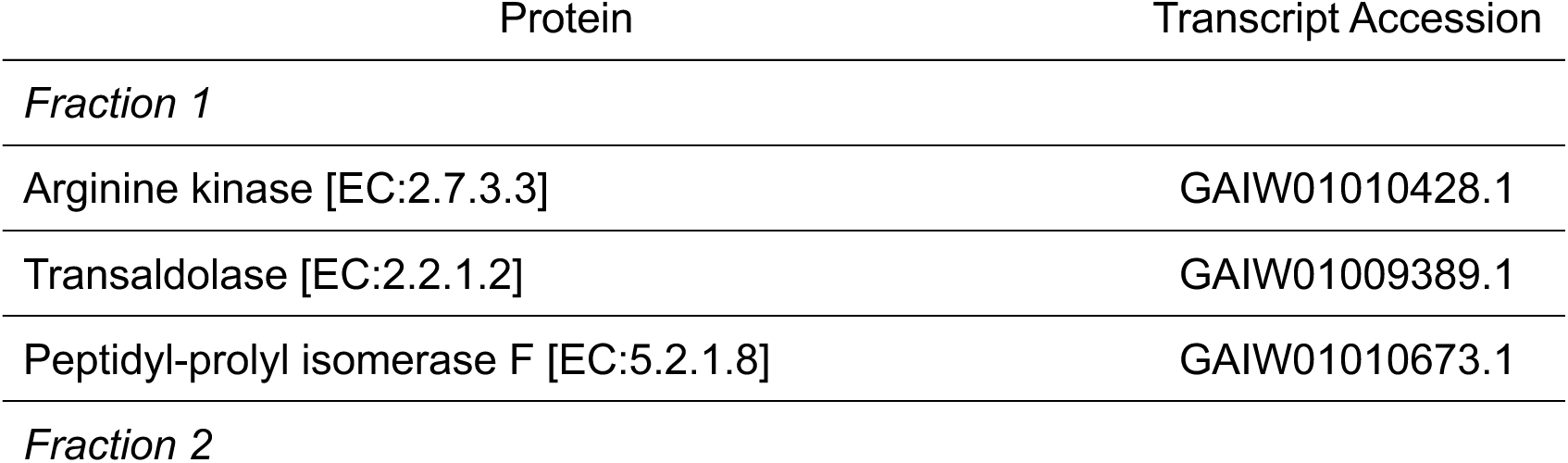
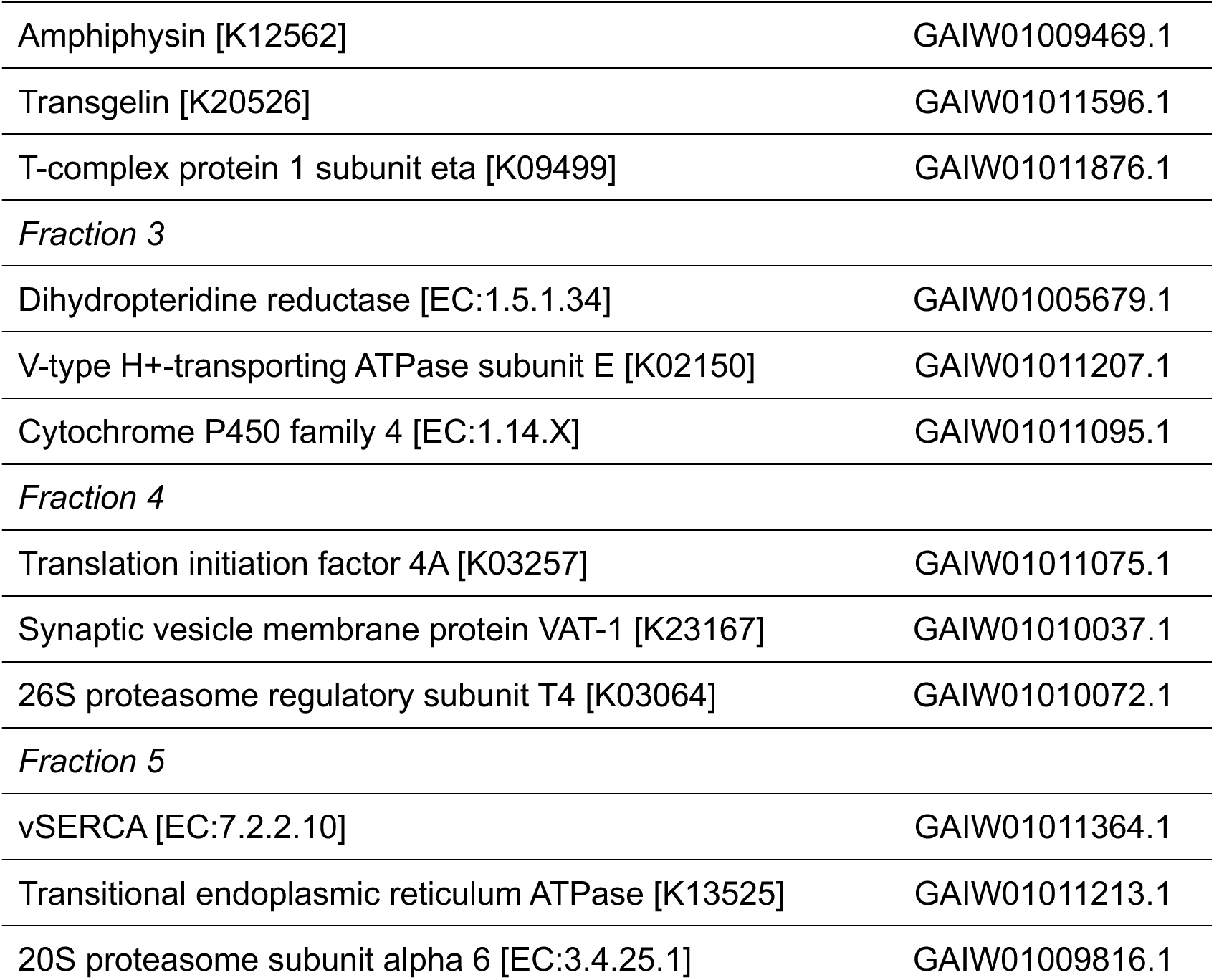
Select proteins identified by mass spectrometry. Protein identity was predicted by Kofam KOALA. Listed proteins are unique to each fraction.

| Protein | Transcript Accession |
| --- | --- |
| *Fraction 1* |  |
| Arginine kinase [EC:2.7.3.3] | GAIW01010428.1 |
| Transaldolase [EC:2.2.1.2] | GAIW01009389.1 |
| Peptidyl-prolyl isomerase F [EC:5.2.1.8] | GAIW01010673.1 |
| *Fraction 2* |  |
| Amphiphysin [K12562] | GAIW01009469.1 |
| Transgelin [K20526] | GAIW01011596.1 |
| T-complex protein 1 subunit eta [K09499] | GAIW01011876.1 |
| *Fraction 3* |  |
| Dihydropteridine reductase [EC:1.5.1.34] | GAIW01005679.1 |
| V-type H<sup>+</sup>-transporting ATPase subunit E [K02150] | GAIW01011207.1 |
| Cytochrome P450 family 4 [EC:1.14.X] | GAIW01011095.1 |
| *Fraction 4* |  |
| Translation initiation factor 4A [K03257] | GAIW01011075.1 |
| Synaptic vesicle membrane protein VAT-1 [K23167] | GAIW01010037.1 |
| 26S proteasome regulatory subunit T4 [K03064] | GAIW01010072.1 |
| *Fraction 5* |  |
| vSERCA [EC:7.2.2.10] | GAIW01011364.1 |
| Transitional endoplasmic reticulum ATPase [K13525] | GAIW01011213.1 |
| 20S proteasome subunit alpha 6 [EC:3.4.25.1] | GAIW01009816.1 |

We used the KofamKOALA algorithm to map our identified venom proteins onto known proteins for functional prediction. Our protein identification results were then compared to the Kyoto Encyclopedia of Genes and Genomes (KEGG) database by the KEGG Mapper tool. This analysis revealed potential functions for each of the venom fractions as represented by membership in KEGG pathway modules (Table 2) (Kanehisa et al., 2017). Most of these putative functions are related to metabolism, including the initial and elongation steps of fatty acid biosynthesis (found in distinct fractions), and glycogen degradation and beta-oxidation, both of which are catabolic processes linked to energy production (Judge and Dodd, 2020).

**Table 2.** KEGG Modules identified among *G. hookeri* venom proteins.

| Function | Proteins | Fraction(s) |
| --- | --- | --- |
| Actin binding proteins | Profilin [K05759]; Cofilin [K05765]; Tropomyosin I [K10373] | 1, 2 |
| Fatty acid biosynthesis, initiation [M00082] | Fatty acid synthase [EC:2.3.1.85], acetyl-CoA carboxylase [EC 6.4.1.2] | 5 |
| Fatty acid biosynthesis elongation [M00083] | Fatty acid synthase [EC:2.3.1.85] | 3, 4 |
| Glycogen degradation [M00855] | Glycogen phosphorylase [EC:2.4.1.1]; Glycogen debranching enzyme [EC:2.4.1.25;3.2.1.33]; Phosphoglucomutase [EC:5.4.2.2] | 4 |
| Beta-oxidation [M00087] | Acyl-CoA dehydrogenase [EC:1.3.8.7]; Acetyl-CoA acyltransferase [EC:2.3.1.16]; Enoyl-CoA hydratase [EC:4.2.1.17] | 4, 5 |

The observed overlap in protein content between fractions (Figure 6A) raises the possibility that vesicles with similar physical properties may also have similar cargo. To test this idea, we analyzed our mass spec data to identify proteins that are specific to F2 and F5 (Table 3) or F3 and F4 (Table 4). F2 and F5 have minimal overlap of protein content, with only 10 proteins in common and 3 proteins that are specific to F2 and F5 and excluded from the other fractions (Table 3). These differences in protein content potentially underlie the observed density difference. On the other hand, F3 and F4 show a great deal of similarity of protein content. In total, 57 proteins are shared between these fractions, and among these we identified 19 proteins that are specifically shared between F3 and F4 and excluded from the other fractions (Table 4). These F3/F4 specific proteins include both predicted metabolic enzymes and proteins predicted to be involved in cell signaling, suggesting that venom vesicle cargos may play multiple functions in host cells.

**Table 3.** Proteins specific to fractions F2/F5 predicted by fraction-specific mass spectrometry.

| Protein | Transcript Accession |
| --- | --- |
| F-type H<sup>+</sup>-transporting ATPase subunit beta [EC:7.1.2.2] | GAIW01012111.1 |
| Superoxide dismutase, Cu-Zn family [EC:1.15.1.1] | GAIW01012119.1 |
| Chitinase [EC:3.2.1.14] | GAIW01012230.1 |

**Table 4.** Proteins specific to fractions F3/F4 predicted by fraction-specific mass spectrometry.

| Protein | Transcript Accession |
| --- | --- |
| Leucine rich repeat protein [COG4886] | GAIW01007069.1 |
| Pyruvate dehydrogenase E1 component subunit alpha [EC:1.2.4.1] | GAIW01010504.1 |
| Reticulon-1 [K20721] | GAIW01010536.1 |
| Alpha-glucosidase [EC:3.2.1.20] | GAIW01010903.1 |
| NADP<sup>+</sup>-dependent farnesol dehydrogenase [EC:1.1.1.216] | GAIW01011046.1 |
| Trehalose 6-phosphate phosphatase [EC:3.1.3.12] | GAIW01011239.1 |
| Presequence protease [EC:3.4.24.-] | GAIW01011585.1 |
| Troponin I [P36188.3] | GAIW01012017.1 |
| Clavesin [K28049] | GAIW01012229.1 |
| Juvenile hormone binding protein [pfam06585] | GAIW01012823.1 |
| Ubiquitin-activating enzyme E1 [EC:6.2.1.45] | GAIW01012873.1 |
| Ezrin-moesin-radixin 1 [K26219] | GAIW01013187.1 |
| Nidogen [K06826] | GAIW01019606.1 |
| Glucose-6-phosphate isomerase [EC:5.3.1.9] | GAIW01021188.1 |
| Aconitate hydratase [EC:4.2.1.3] | GAIW01021311.1 |
| Transcriptional activator protein Pur-alpha [K21772] | GAIW01021479.1 |
| Dipeptidyl-peptidase III [EC:3.4.14.4] | GAIW01021615.1 |
| Unannotated | GAIW01022664.1 |
| Unannotated | GAIW01023502.1 |

### Venom vesicles gain entry into host plasmatocytes

We hypothesized that venom vesicles may serve as venom protein transport mechanisms. If this is the case, we would predict that venom vesicles would interact with and potentially enter into, host cells. To assess this, we stained whole venom with ExoSparkler (Exo-Venom), a lipophilic dye that enables tracking of vesicle internalization by fluorescence microscopy (Shimomura et al., 2020). PBS alone was incubated with ExoSparkler (Exo-PBS) to control for any background staining or dye carryover following dye removal. Exo-Venom and Exo-PBS were then incubated with primary hemocyte cultures established from *hop[Tum]* third instar larvae. *hop[Tum]* is a gain of function mutation in the JAK-STAT pathway which leads to the ectopic production of lamellocytes (Hanratty and Dearolf, 1993), allowing us to assess vesicle interactions with both plasmatocytes and lamellocytes. After 40 minutes of incubation with Exo-Venom or Exo- PBS, cells were fixed and imaged to assess ExoSparkler localization (Figure 7A-D; Figure S3).

**Figure 7.**
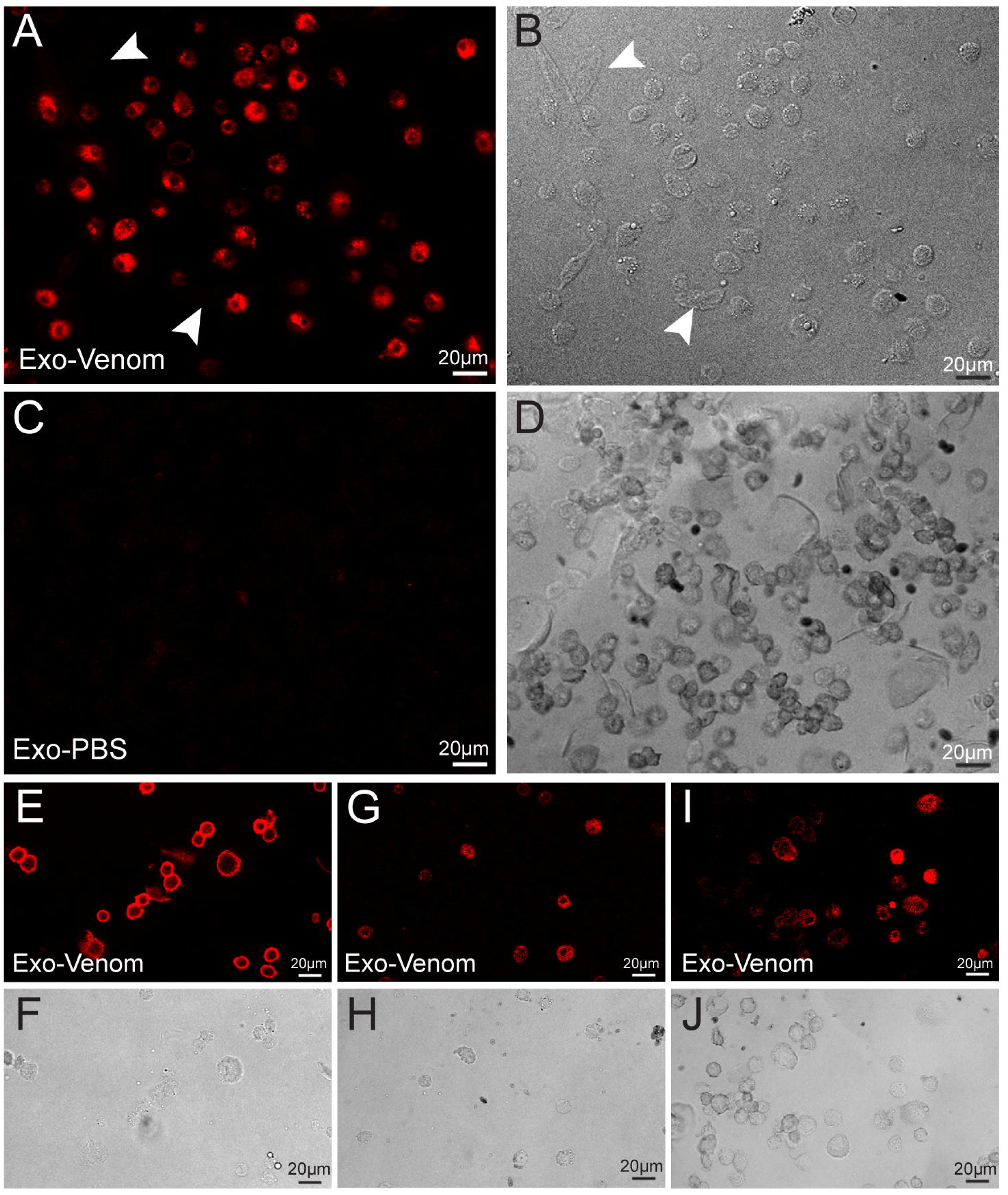
Analysis of venom vesicle entry into host cells. ExoSparkler labeled venom vesicles (Exo-Venom) compared to ExoSparkler control (Exo-PBS). Scale bars indicate 20 µm in all panels. (A-D) *hop[Tum]* hemocyte primary cultures incubated with Exo- Venom (A-B) and Exo-PBS (C-D). Confocal micrographs (A, C) shown with paired brightfield images (B, D). Arrowheads in (A-B) indicate unstained lamellocytes. (E-J) Time course of *w[1118]* hemocyte primary cultures. For time course analysis, cells were fixed following 5 minutes (E-F), 15 minutes (G-H), or 40 minutes (I-J) of incubation with Exo- Venom. Confocal micrographs (E, G, I) shown with paired brightfield images (F, H, J).

We observed that Exo-Venom incubated plasmatocytes had high levels of fluorescence, both at the membrane and within the cytoplasm (Figure 7A-B; Figure S3A- B), although the signal appears to be excluded from the nucleus (Figure 7A; Figure S3A). We did not observe detectable levels of fluorescence in lamellocytes (Figure 7A-B, indicated by arrowheads), suggesting that *G. hookeri* vesicles may specifically target plasmatocytes. Fluorescence was largely undetectable in Exo-PBS treated hemocytes (Figure 7C-D; Figure S3C-D), suggesting that our dye cleanup was successful and that the fluorescence entry into host cells is specific to stained venom vesicles.

The apparent localization of ExoSparkler within the cytoplasm of plasmatocytes suggested that vesicles may be gaining entry into host cells. To better understand this process, we performed a time course experiment to track the interaction. We incubated primary plasmatocyte cultures from *w[1118]* larvae with Exo-Venom for 5, 15, and 40 minutes before fixation and imaging. We observed that at the earliest time point, fluorescence signal was localized nearly exclusively to the plasma membrane (Figure 7E- F). After 15 minutes of incubation, cytoplasmic fluorescence was observed (Figure 7G- H), and by 40 minutes nearly all of the signal was internalized into the cytoplasm (Figure 7I-J). These findings suggest that venom vesicles gain entry into host plasmatocytes, and that this process is mediated by a stepwise mechanism.

## Discussion

To test our hypothesis that *G. hookeri* venom proteins, such as the large hydrophobic venom SERCA, are transported into host cells via venom vesicles, we characterized the composition of *G. hookeri* venom through biophysical, cell biological, and biochemical approaches. Our findings reveal that these venom proteins are associated with a heterogenous population of lipid-bound small EVs, which appear to facilitate their entry into *D. melanogaster* immune cells. It seems likely that at least one of these vesicle types serves to transport venom SERCA into the host, and our finding that vesicle entry targets host plasmatocytes and largely excludes lamellocytes in culture aligns with our previous findings that *G. hookeri* venom SERCA activity is specific to plasmatocytes during infection (Mortimer et al., 2013).

Our biophysical characterization of venom vesicles highlights potential differences in vesicle usage between *G. hookeri* and other Figitidae with described venom vesicles (Table 5). *L. heterotoma* extracellular vesicles, formerly known as virus-like particles, compose a generally homogenous population of large (300 nm) vesicles with a distinct knob-like morphology (Morales et al., 2005; Chiu et al., 2006). The density of these extracellular vesicles has not been determined, however they are purified as a single fraction from a Nycodenz gradient, suggesting that the density is likewise uniform (Heavner et al., 2017). The venom vesicles from *L. boulardi*, commonly known as venosomes, are 100 - 200 nm in diameter, and are also found in an aggregated state with aggregates having a total size of 400 nm - 1µm (Wan et al., 2019). There is disagreement over the morphology of these venosomes, with reports differing between a stellate or round morphology (Gueguen et al., 2011; Wan et al., 2019). It is unclear from previous reports whether these morphological differences represent unique vesicle subpopulations within *L. boulardi* venom or if they may be artefacts of sample preparation or imaging. Again, although the density has not been determined, these venosomes are purified as a single fraction at 25-30% Nycodenz (Wan et al., 2019). In contrast, *G. hookeri* have as many as 6 separable populations of venom vesicles (Table 5). This includes low-density small EVs (F2 vesicles), and two populations of medium-density small EVs, that are separable by zeta potential (F3 and F4b vesicles with mean zeta potentials between -12 mV and -21 mV, and F4a vesicles with a mean zeta potential of -65 mV). Finally, we have identified two populations of high-density EVs that otherwise differ by size, morphology, and zeta potential (F5a and F5b vesicles, Table 5).

**Table 5.**
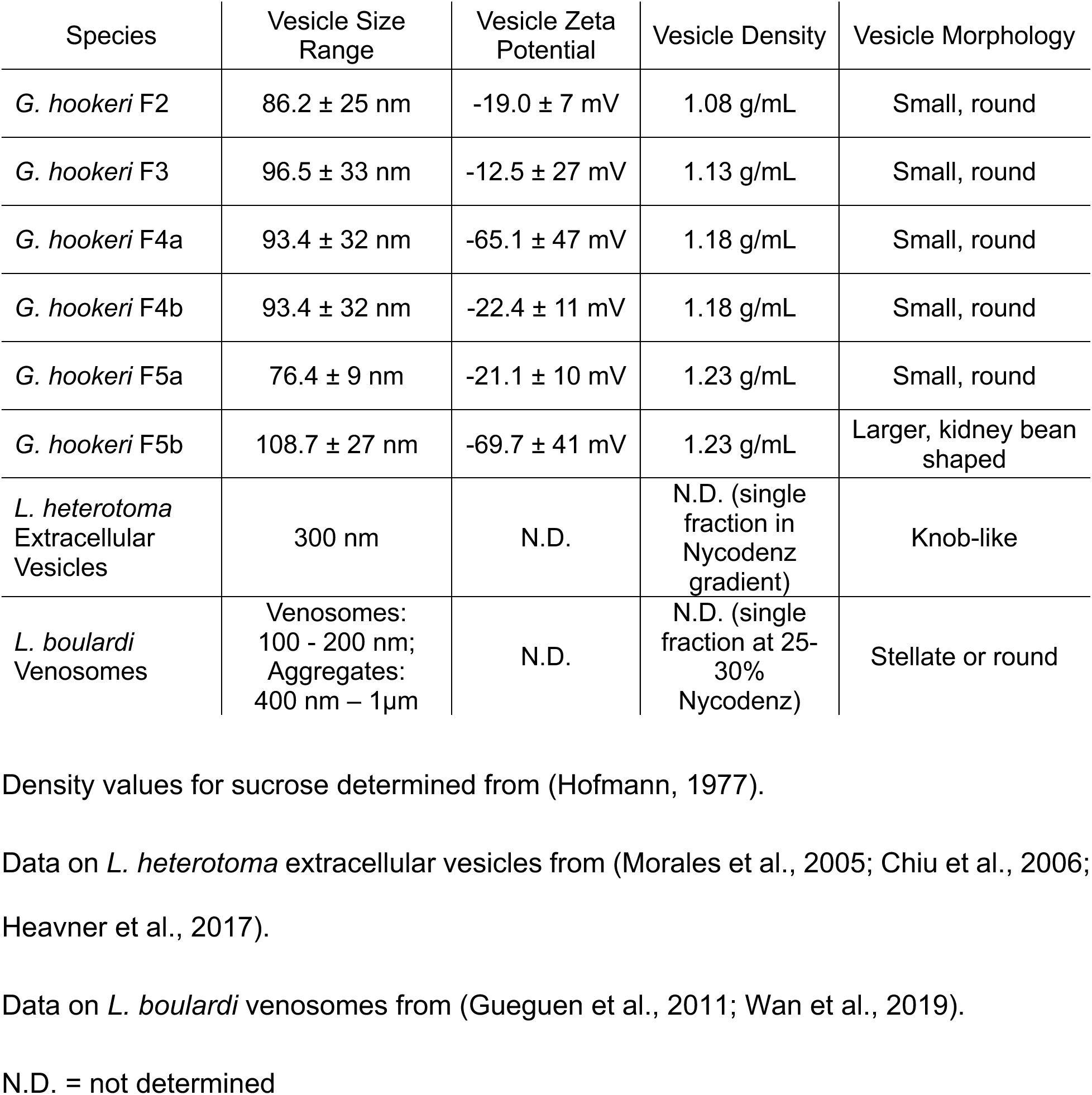
Comparison of venom vesicle morphology and biophysical characterization with *G. hookeri* venom and across parasitoid species.

It is important to note that methodology differences between TRPS and NTA (electrical vs optical measurements) can account for the slight differences in vesicle diameter that we measured between experiments (Anderson et al., 2013; Vogel et al., 2021). However, the relative differences between fractions are very consistent between these disparate methods, and our TRPS data match reasonably well with sizes estimated from TEM imaging, suggesting that our presented vesicle diameter measurements likely provide an accurate reflection of the *G. hookeri* venom composition.

To begin characterizing vesicle cargo, we performed fraction-specific mass spectrometry, and identified unique proteins in each fraction, along with proteins found across multiple fractions. Similar to previous reports (Heavner et al., 2013, 2017) we discovered additional venom proteins with the optimization of the protein purification approach to recover vesicle proteins in comparison with previous venom mass spectrometry that assumed soluble proteins and complexes (Mortimer et al., 2013). Our mass spectrometry results suggest that vesicle cargos likely contain venom proteins with a wide range of possible functions. We further hypothesize that each vesicle class we have described plays a distinct functional role in infection. In order to test this hypothesis and more precisely determine vesicle cargos, we will need to further optimize our vesicle purification approach.

Our combined results from mass spectrometry and SDS-PAGE suggest that the more dense fractions contain larger proteins, although in both cases, we discovered a large range of protein sizes in each fraction. Because density is not merely a product of size, this suggests that larger proteins may be more likely to be found associated with higher density particles, such as vesicles. Similarly, while there are differences in the mean protein isoelectric point across fractions, these differences are not predictive of the vesicle zeta potentials measured. Because zeta potential reflects the surface potential of the vesicle, it is likely driven primarily by lipid composition with contributions from the proteins found on the surface of the vesicle membrane (Midekessa et al., 2020; Ghadami and Dellinger, 2023). Additionally, our data show a small but significant negative correlation between vesicle size and zeta potential, likely revealing a relationship between these properties that is also strongly influenced by the other factors discussed.

More generally, our findings suggest that the characterization of newly discovered vesicle populations should include more than EM analysis or size determination in isolation (Table 5). We uncovered morphologically similar vesicles that are distinct when compared on other parameters. For instance, F2 vesicles and F5a vesicles have similar sizes, morphology, and zeta potential, but display dramatically different densities and have minimal overlap in protein content. Likewise, although F4 vesicles appear homogeneous in terms of morphology, size, and density, F4 contains two subpopulations of vesicles with drastically different zeta potentials, suggesting the potential presence of different functional populations of vesicles within this fraction.

Our characterization of *G. hookeri* venom vesicles provides insight into the intricate virulence strategies used by parasitoid wasps to overcome host defenses, and may lead to further discoveries in the principles underlying protein packaging and transport in membrane enclosed vesicles.

## Experimental procedures

### Insects

Flies were maintained on standard Drosophila medium of cornmeal, yeast, and molasses. The wild type controls *w[1118]* (RRID: BDSC_5905) and *Canton-S* (RRID: BDSC_64349), and the *hop[Tum]* (RRID: BDSC_8492) mutant *Drosophila melanogaster* strains were obtained from the Bloomington Drosophila Stock Center (Bloomington, IN) (RRID: SCR_006457). The *Drosophila yakuba* strain Tai18E2 (RRID: NCBITaxon_7245) was obtained from Kyorin-Fly (Tokyo, Japan). *Ganaspis hookeri* (NCBI: txid2777979) wasps were maintained on *D. yakuba* as previously described (Mortimer et al., 2013).

### Venom purification

To extract *G. hookeri* venom, the venom apparatus was dissected from adult female wasps into HALT™ protease inhibitor (Thermo Fisher Scientific) diluted to 1X in phosphate buffered saline (PBS: 137 mM NaCl, 2.7 mM KCl, 10 mM Na2HPO4, 1.8 mM KH2PO4, pH 7.4). Dissected tissues were pelleted by centrifugation at 8,000 x g and homogenized (Pestle Homogenizer, Fisher Scientific) in short pulses of 2 seconds under non-lysing conditions. The homogenized sample was incubated on ice for 5 minutes and centrifuged at 1,000 x g for 10 minutes to pellet cells, cellular debris, and any remaining tissues. Following centrifugation, the supernatant was removed from the pellet and stored at 4**°**C. We observed venom vesicle aggregation at 4°C, so venom samples were filtered using a 0.22 µm PES syringe filter (SFPES004022N, Membrane Solutions) prior to use. SDS-PAGE analysis of fresh and filtered venom samples demonstrates that venom composition is unchanged by filtration (Figure S4). See (Bretz and Mortimer, 2025a) for a step-by-step protocol.

### Detergent treatment for vesicle disruption

To test whether venom vesicles are sensitive to detergent disruption, purified venom from 30 *G. hookeri* wasps was incubated with 0.1% SDS (w/v), 1% SDS (w/v), or PBS control for one hour before further analysis as described below.

### Fluorescent vesicle labeling

To label *G. hookeri* venom vesicles, 100 µL of 50 purified venom was combined with 2 µL of ExoSparkler Membrane Labeling dye (Dojindo), resuspended at 1X in 10 µL dimethyl sulfoxide (DMSO, Fisher Scientific), and incubated at 37°C for 30 minutes. A control sample of 100 µL PBS was simultaneously incubated with 2 µL of ExoSparkler at 37°C for 30 minutes. Both venom and control samples were then transferred to the provided ExoSparkler collection tubes and centrifuged at 3,000 x g for 5 minutes at room temperature for the removal of unbound dye. Samples were washed twice with PBS and spun dry at 3,000 x g. Labeled venom and control samples were recovered in 50 µL of PBS.

### Confocal microscopy

Confocal microscopy was performed on a Leica SP8 confocal microscope (Leica Microsystems) using a HC PL APO 40x/1.10 W motCORR CS2 objective or a HC PL APO 63x/1.40 OIL CS2 objective as indicated in the text, and a white light laser set to the appropriate excitation wavelengths in a room temperature of 19°C.

### Whole mount venom apparatus imaging

For whole mount imaging, the venom apparatus was dissected from 10 adult female wasps into PBS. Tissues were fixed in 4% paraformaldehyde in PBS (w/v, Fisher Scientific) for 10 minutes, and permeabilized in PBS + 0.1% Triton-X100 (PBSTx) for 10 minutes. Tissues were then blocked in 5% normal goat serum in PBSTx and stained with the α-SERCA 809-27 antibody (Moller et al., 1997) diluted 1:1,000 in blocking solution, and FITC-conjugated fluorescent secondary antibody diluted 1:500 in blocking solution (Jackson ImmunoResearch). Venom apparatuses were co-stained with 165nM phalloidin-Alexa Fluor 633 (Molecular Probes) in PBS for 10 minutes. Stained tissues were mounted whole in VectaShield (VectorLabs) and imaged on the SP8 confocal as described above. Z-stack images spanning the depth of the tissue were taken at 40x and 63x magnification as indicated in the text. LAS X Navigator was used to define regions of interest and stitch Z-stack images taken in multiple fields of view. Z-stacks were processed in FIJI (Schindelin et al., 2012) for maximum intensity 2D projections, and “Brightest Point” 3D projections using default parameters.

### Fluorescently labeled vesicle imaging

To assess venom vesicle entry into *D. melanogaster* plasmatocytes, we incubated venom vesicles fluorescently labeled with ExoSparkler (see above) with primary hemocyte cultures and used confocal microscopy to image vesicle entry. Primary hemocyte cultures were derived from second instar *hop[Tum]* larvae: larvae were rinsed in PBS, and bled in 35 µL of PBS on a dissection slide (Tekdon, Inc) in duplicate. Cells were allowed to become adherent for 10 minutes. PBS was removed and 30 µL of ExoSparkler-venom or ExoSparkler-PBS control were added, and cells were incubated for 40 minutes shielded from light. Solutions were removed and cells were fixed with 4% paraformaldehyde in PBS for 15 minutes (Fisher Scientific), washed in PBS and mounted in VectaShield (VectorLabs). Cells were imaged on the SP8 confocal at 40x magnification as described above.

### Vesicle entry time course imaging

To investigate the process of venom vesicles entry into host plasmatocytes, primary plasmatocyte cultures were derived from second instar *w[1118]* larvae as described above. Cells were incubated with 30 µL ExoSparkler-venom for 5, 15, or 40 minutes, then fixed in 4% paraformaldehyde in PBS for 15 minutes. Cells were mounted in VectaShield (VectorLabs), and imaged on the SP8 confocal at 40x magnification as described above.

### Sucrose gradient fractionation

To separate *G. hookeri* venom proteins and vesicles by density, purified venom was subjected to sucrose gradient fractionation through a discontinuous gradient ranging from 10 - 50% (w/v) sucrose. A PBS buffered 66% (w/v) sucrose solution was diluted in PBS to final concentrations of 50%, 40%, 30%, 20%, and 10% on ice. The discontinuous gradient was generated by underlaying progressively more dense solutions in thick-wall polycarbonate tubes (11x34 mm, Beckman Coulter). 100 µL of venom or PBS only control were applied to the top of the sucrose gradient and samples were centrifuged in a swinging bucket rotor (TLS-55, Beckman Coulter) at 120,000 x g for 20 hours at 4°C (Beckman Optima TLX) without braking.

### Sodium Dodecyl Sulfate-Polyacrylamide Gel Electrophoresis (SDS-PAGE)

SDS-PAGE was used to analyze venom protein profiles from sucrose gradient fractionation. Purified venom from 60 female wasps was fractionated by sucrose gradient fractionation protocol. Fractions were dialyzed into 100 µL of PBS using 2k MWCO Slide-A-Lyzer (ThermoFisher) dialysis cups with two room temperature buffer exchanges and an overnight dialysis at 4°C. For SDS-PAGE analysis, 20 µL of venom from each fraction was sonicated at full power using a Branson CPX1800H ultrasonic water bath (Branson Ultrasonics) for 15 minutes, and combined with 4 µL of 6X Laemmli SDS sample buffer (Thermo Scientific). Samples were heated at 95°C for 5 minutes. The entire sample volume was loaded on a 4–20% Mini-PROTEAN TGX Precast Protein Gel (Bio-Rad) in 1X Tris/glycine/SDS buffer and run at 40 mA constant until the dye front approached the end of the gel. The PAGE gel was removed from the electrophoresis rig and stained for one hour using GelCode Blue Stain Reagent (Thermo Scientific).

### Western blotting

For Western blotting, sample preparation and SDS-PAGE were performed as described above. Immobilon PVDF membrane (Millipore Sigma) was prepared by activation in 100% methanol for 1 minute, a 2-minute rinse in DI water, then a 10-minute incubation in Towbin transfer buffer (25 mM Tris, 192 mM glycine, 20% (v/v) methanol). The blotting sandwich was assembled and placed in the Trans-Blot Turbo Transfer System (Bio-Rad) for 30 minutes at 1.0 A maximum, 25V constant. The blot with transferred protein was removed and blocked at room temperature for 30 minutes with EveryBlot Blocking Buffer (Bio-Rad). The blot was removed and incubated with α-SERCA primary antibody (sc-271669, Santa Cruz Biotechnology) at a 1:500 dilution in 6 mL total volume of 1:1 TBS-T (20 mM Tris, 150 mM NaCl, 0.1% Tween 20 (w/v, Sigma-Aldrich)): EveryBlot Blocking Buffer overnight at 4°C. Primary antibody solution was removed, washed twice in 10 minute increments with TBS-T, and a final rinse in TBS. IRDye 800CW Goat anti-Mouse IgG secondary antibody (Licor) was diluted 1:15,000 in a TBS-T:Block solution and was incubated for 1 hour at room temperature on the blot. Following the previously described wash steps, the blot was shielded from light and dried at room temperature, then imaged using the Odyssey CLx imager (LICORbio).

### Electron microscopy

To examine venom composition and vesicle morphology, purified venom from 50 female wasps was prepared and imaged using electron microscopy techniques. For scanning electron microscopy (SEM), venom samples were filtered, fixed in 5% glutaraldehyde in 2X PBS, and imaged on a Helios NanoLab 650 Dual Beam FEG SEM (FEI/Thermo Fisher, Hillsboro, OR). For transmission electron microscopy (TEM), venom samples were filtered, and either fixed (whole venom, fractions F2-F4) in 5% glutaraldehyde in 2X PBS, or left unfixed (fraction F5). A drop of sample was placed on a glow-discharged grid, which was then briefly rinsed in a bead of water and stained with uranyl acetate. Excess stain was wicked off and grids were imaged on a Titan G2 ChemiSTEM TEM (FEI/Thermo Fisher, Hillsboro, OR).

### Nanoparticle tracking analysis data collection

Venom samples were subjected to sucrose gradient fractionation, and five fractions were collected to isolate the venom vesicles. The collected fractions were then dialyzed against PBS to remove sucrose. Immediately prior to performing nanoparticle tracking analysis (NTA), dialyzed fractions were filtered using a 0.22 µm PES syringe filter and diluted 5-fold with PBS. The hydrodynamic diameters of vesicles were measured using a NanoSight LM10, configured with a temperature controlled LM14 sample viewing unit equipped with a 532 nm laser and a high sensitivity sCMOS camera (Malvern Panalytical). The temperature of the sample was held constant at 22°C, and analysis was performed under a constant flow rate of 10 μL/min to ensure representative sampling. NanoSight 3.1 software was used to collect five 60 s videos of each sample with the camera level set to 14. Using a detection threshold of 2, the mean hydrodynamic diameter and standard deviation of the vesicles in each sample were obtained from the analysis of approximately 10,000-55,000 individual vesicles, depending on the vesicle concentration in the collected fraction.

### Tunable resistive pulse sensing

For the assessment of venom vesicle characteristics, tunable resistive pulse sensing (TRPS) was performed using the Exoid (Izon). Venom was purified from 75 *G*. *hookeri* females and dialyzed in PBS in 2k MWCO Slide-A-Lyzer (ThermoFisher) dialysis cups with two 2-hour buffer exchanges at room temperature, and a final exchange overnight at 4°C in fresh PBS. Following dialysis, venom samples were filtered with a 0.22 um PES filter (SFPES004022N, Membrane Solutions). Samples were analyzed using an NP100 Nanopore (A97068, Izon) or NP250 Nanopore (A88596, Izon). Solutions were prepared according to manufacturer’s instructions with the following exception: for TPRS analysis of detergent treated venom, the SDS concentration of the measurement electrolyte was adjusted to equal the SDS concentration of the sample.

For size and concentration measurements of whole and fractionated venom, the NP100 Nanopore was calibrated with CPC 100 calibration particles and set to a stretch of 45 mm. For measurements of detergent treated venom, samples were measured using an NP250 nanopore at a 47 mm stretch. Calibration was performed utilizing CPC200 nanoparticles. Measurements were taken at three pressures: P1: 1000 Pa, P2: 800 Pa, and P3:1400 Pa, while a constant Nanopore stretch and constant 1100mV voltage were maintained throughout the measurement session. A minimum of 500 particles were counted at each pressure; the particle measurement rate never exceeded 880 particles / minute, and the root mean square (RMS) noise never exceeded 12 pA to ensure measurement accuracy.

For zeta potential measurements, the NP100 Nanopore was calibrated with CPC 100 calibration particles and set to a stretch of 45 mm. Calibration analysis was performed at variable voltages and applied pressures (V1P1: 766 mV, 100 Pa; V1P0: 766 mV, 0 Pa; V2P0: 613 mV, 0 Pa; V3P0: 460 mV, 0 Pa). Sample measurements were taken at a constant Nanopore stretch of 45mm and constant 766 mV voltage, without any applied pressure (0 Pa). The average particle measurement rate was 29.3 particles / minute.

Data were exported as comma-separated values (CSV) files using the Exoid Control Suite software (version 1.4). See (Bretz and Mortimer, 2025b) for a step-by-step protocol for TRPS measurements.

### Sample Preparation for MS Analysis

35 µL of each sucrose gradient fraction in SDS sample buffer (2% SDS,10% glycerol, 50 mM Tris, 0.5X PBS, trace bromophenol blue) were separated onto a NuPAGE™ 10% Bis-Tris Protein gel (Cat. # NP0315BOX, Invitrogen) at 200V constant for 10min. The gel was stained using a Colloidal Blue Staining Kit (Cat. #LC6025, Invitrogen) following the manufacturer’s instruction. Each gel lane was excised and digested overnight at 37°C with Pierce™ Trypsin Protease, MS Grade (Cat. #90058, Thermo Scientific) as per manufacturer’s instruction. Digests were reconstituted in 0.1%FA in 5:95 ACN: ddH2O at ∼0.1 µg/µL.

### nLC-ESI-MS2 Analysis and Database Searches

Peptide digests (8 µL each) were injected onto a 1260 Infinity nHPLC stack (Agilent Technologies), and separated using a 100 µm inner diameter x 12.5 cm pulled tip C-18 column (JupiterC-18 300 Å, 5 microns, Phenomenex). This system runs in-line with a Thermo Orbitrap Velos Pro-hybrid mass spectrometer, equipped with a nano-electrospray source (Thermo Fisher Scientific), and all data were collected in CID mode. The nHPLC was configured with binary mobile phases that included solvent A (0.1% FA in ddH2O), and solvent B (0.1% FA in 15%ddH2O / 85% ACN), programmed as follows; 10min @ 5%B (2µL/ min, load), 90min @ 5% -40%B (linear: 0.5nL/ min, analyze), 5min @ 70%B (2µL/ min, wash), 10min @ 0%B (2µL/min, equilibrate). Following each parent ion scan (300 -1200m/z @ 60k resolution), fragmentation data (MS2) was collected on the topmost intense 15 ions. For data-dependent scans, charge state screening and dynamic exclusion were enabled with a repeat count of 2, repeat duration of 30s, and exclusion duration of 90s.

The XCalibur RAW files were collected in profile mode, centroided and converted to MzXML using ReAdW v. 3.5.1. The data were searched using G1 protein sequences and SEQUEST, which was set for two maximum missed cleavages, a precursor mass window of20ppm, trypsin digestion, variable modification C @ 57.0293, and M @ 15.9949.

### Peptide Filtering, Grouping, and Quantification

Peptides identified by mass spectrometry were mapped back onto the *G. hookeri* transcriptome (Sequence Read Archive: SRR805628) by SEQUEST (Thermo Fisher Scientific) search. SEQUEST results were filtered using Scaffold (Protein Sciences, Portland Oregon). Scaffold filters and groups all peptides to generate and retain only high confidence IDs while also generating normalized spectral counts (N-SC’s) across all samples for the purpose of relative quantification. The filter cut-off values were set with minimum peptide length of >5 AA’s, with no MH+1charge states, with peptide probabilities of >80% C.I., and an FDR <1.0%. Scaffold incorporates the two most common methods for statistical validation of large proteome datasets, the false discovery rate (FDR) and protein probability. Relative quantification across experiments was then performed via spectral counting, and when relevant, spectral count abundances were then normalized between samples.

### Bioinformatics analysis of identified venom proteins

Fraction membership was determined according to the following criteria: First, a protein must be identified by at least 2 unique peptides to count as present in a fraction. Second, the protein must have been identified in the whole venom control sample, or have a read coverage of at least 10 total peptides. Third, the total spectra count must be higher within the fraction than the average total spectra count across the fractions. These criteria allowed us to focus on proteins that have a high likelihood of presence in a given fraction and decreasing noise from any potential carryover between fractions.

To determine protein identities, we used the KofamKOALA webserver (https://www.genome.jp/tools/kofamkoala/; accessed 04-2025) to map our positively identified venom proteins onto established KEGG Orthology (KO) groups (Aramaki et al., 2020). The KO assignments were additionally used for functional annotation using the KEGG Mapper Reconstruct tool (https://www.kegg.jp/kegg/mapper/reconstruct.html; accessed 04-2025) to identify protein “modules”, groups of proteins which form complete signaling or metabolic pathways (Kanehisa and Sato, 2020; Kanehisa et al., 2022). We used FlyBase (release FB2025_02) to find information on *D. melanogaster* phenotypes and the integrated BLAST tool to map *G. hookeri* proteins onto *D. melanogaster* homologs (Öztürk-Çolak et al., 2024).

### Statistical analysis

Statistical analyses were performed in R (RRID: SCR_001905) (R Core Team, 2021), and data were plotted using the ggplot2 (RRID: SCR_014601) and ggpubr (RRID: SCR_021139) packages (Wickham, 2009; Kassambara, 2020).

Mixture modeling of diameter and zeta potential data from TRPS was performed using the mixtools package (Benaglia et al., 2010). Recombined fractions were analyzed to reflect whole venom properties. Each model distribution is characterized by μ (population mean), σ (standard deviation) and λ (weight, or proportion of the total data points attributed to the distribution). We used a λ threshold of 10 (i.e. 10% of the vesicles in the population) to accept a distribution, and the resulting distributions were plotted by the plot_mix_comps command from plotGMM (Waggoner and Chan, 2020). For fractions where diameter or zeta potential appeared monomodal, the summarySE function from Rmisc (Hope, 2022) was used to calculate the mean and standard deviation, which were overlaid on each histogram to summarize the spread of the data.

For the detergent sensitivity experiment, the loess (local weighted regression) line was calculated from the size vs concentration scatterplot in ggpubr. Linear regression lines were calculated in ggpubr for size vs zeta potential scatterplots. Pearson’s correlation between size and zeta potential was additionally calculated in ggpubr.

Molecular weight and pI of proteins identified by mass spectrometry, were compared by one-way ANOVA followed by Tukey’s HSD *post hoc* testing for pairwise comparisons using multcomp (RRID: SCR_018255) (Hothorn et al., 2008). The Venn diagram describing protein fraction membership was created using ggVennDiagram (Gao et al., 2021).

## Supporting information

This article contains supporting information.

## Acknowledgments

We would like to thank Carrie Marean-Reardon, Kristen Snitchler, Michael Youkhateh, and other members of the Venom Biochemistry and Molecular Biology Lab for their assistance and feedback. We would also like to thank Dr. Kevin Edwards for assistance with confocal microscopy, and Drs. Sarah Clark, Evan Forsythe, Kenton Hokanson, Colin Johnson, and Benjamin Philmus for helpful discussions of the project. *Drosophila melanogaster* lines obtained from the Bloomington Drosophila Stock Center (NIH P40OD018537) were used in this study. The ISU Confocal Microscopy Facility was funded by NSF grant DBI-1828136.

## Funding and additional information

This work was supported by NIH grant R35 GM133760 to NTM, the Linus Pauling Institute Harvey H & Donna Morre Graduate Research fellowship to NMB, and start-up funds from Oregon State University and Illinois State University to NTM and JDD. The content is solely the responsibility of the authors and does not necessarily represent the official views of the National Institutes of Health.

