## Supplementary figures and images for "Venom vesicles from the parasitoid *Ganaspis hookeri* facilitate venom protein entry into host immune cells"

### Figure S1

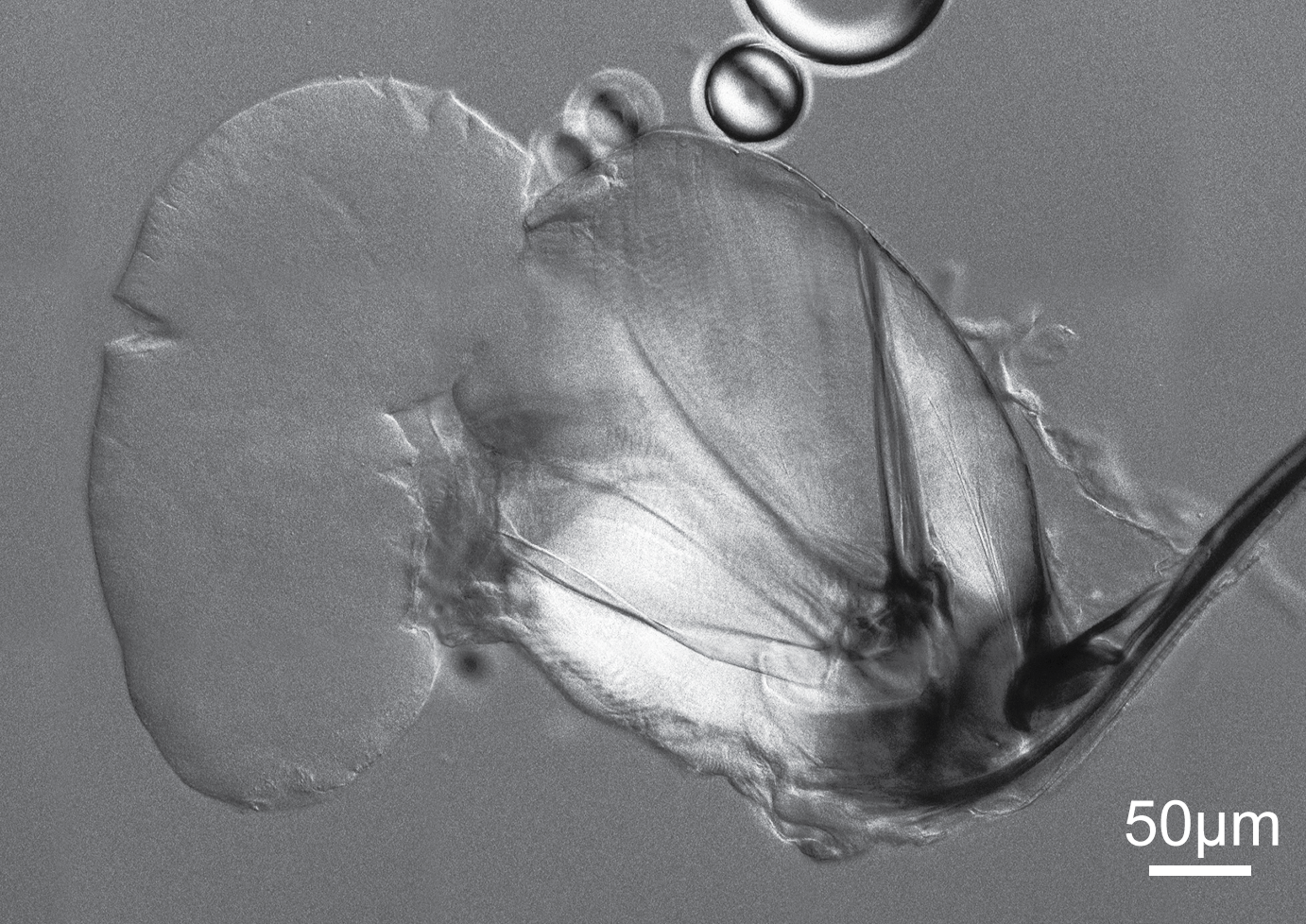

### Figure S2

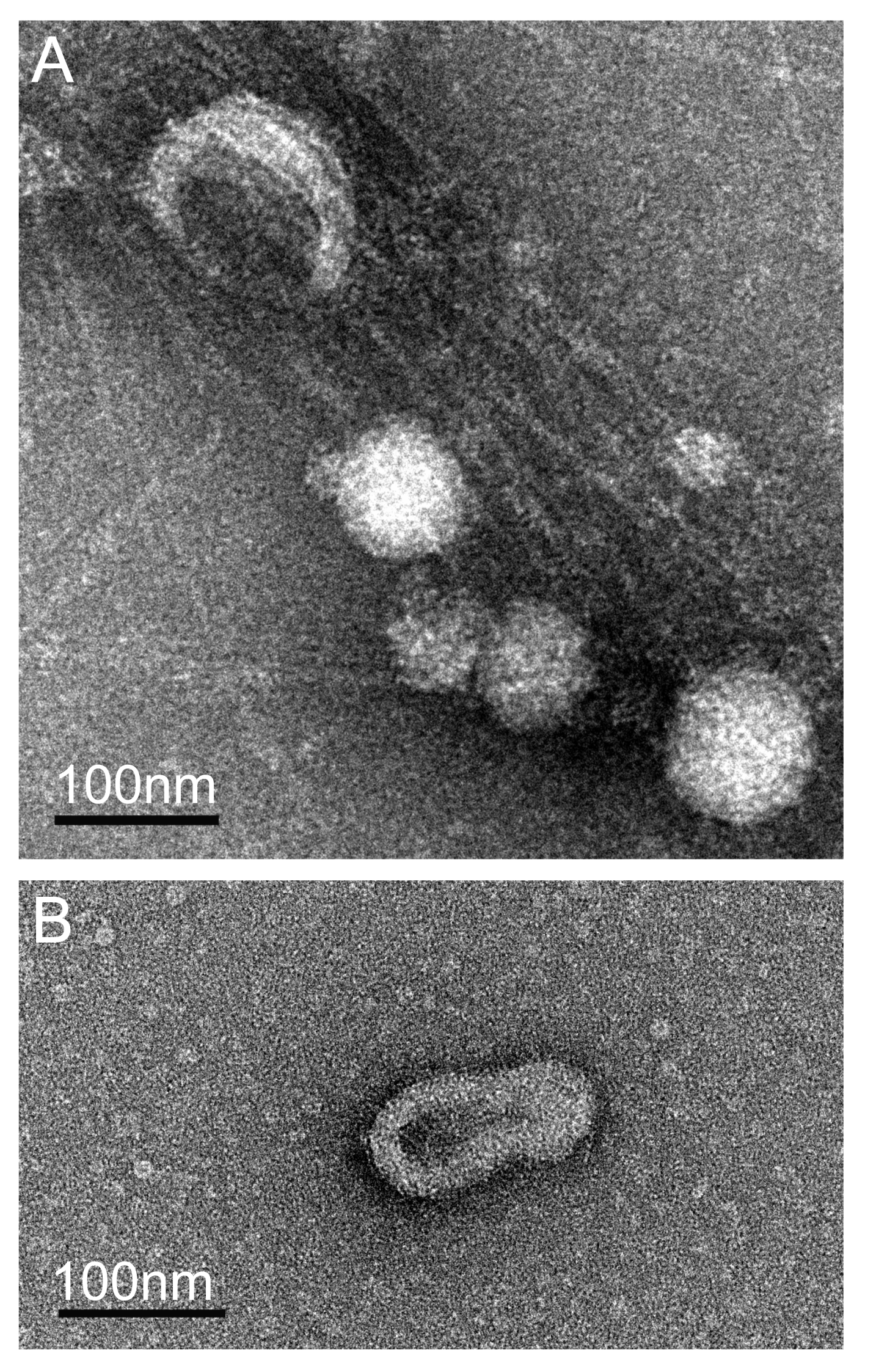

### Figure S3

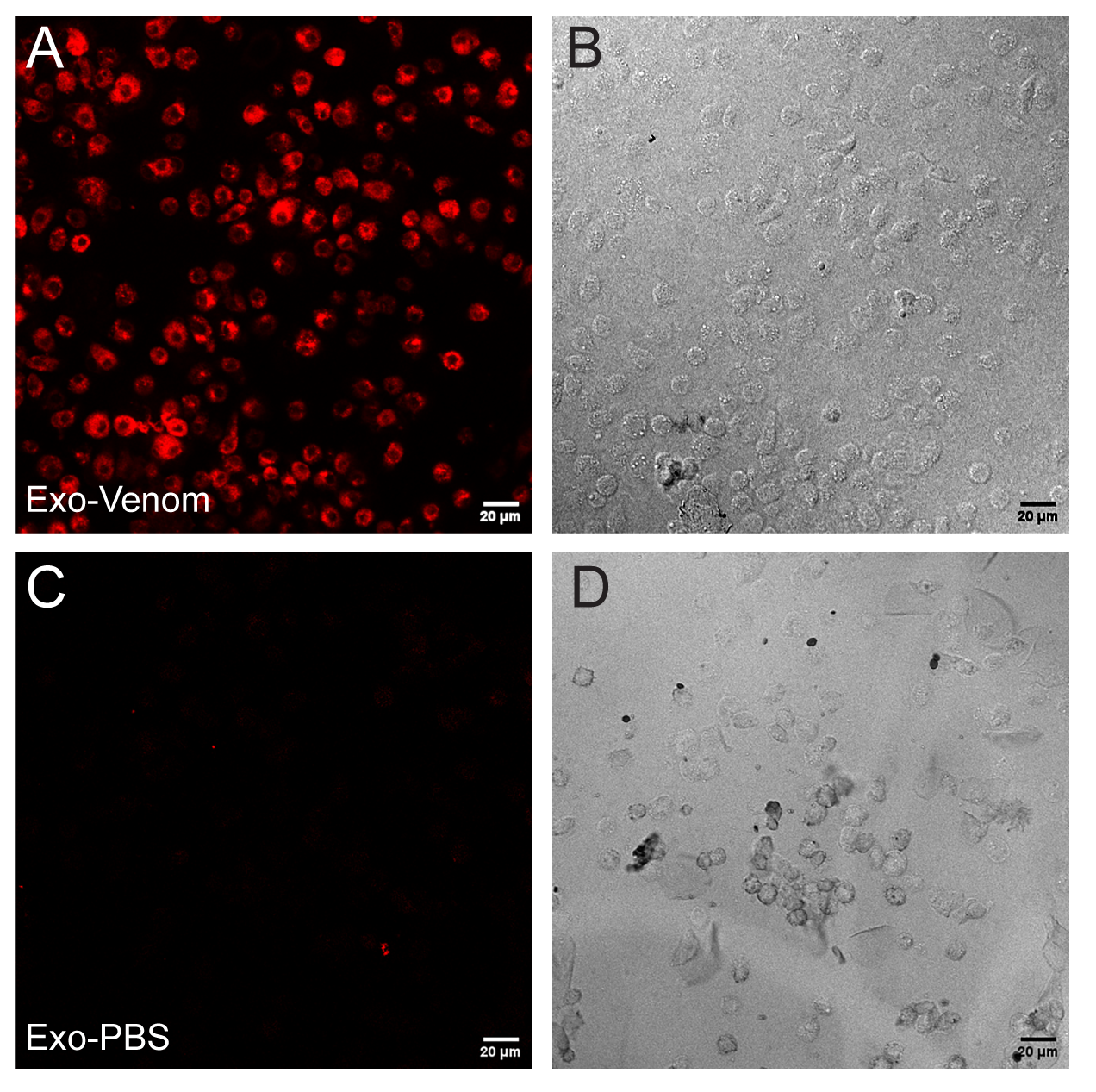

### Figure S4

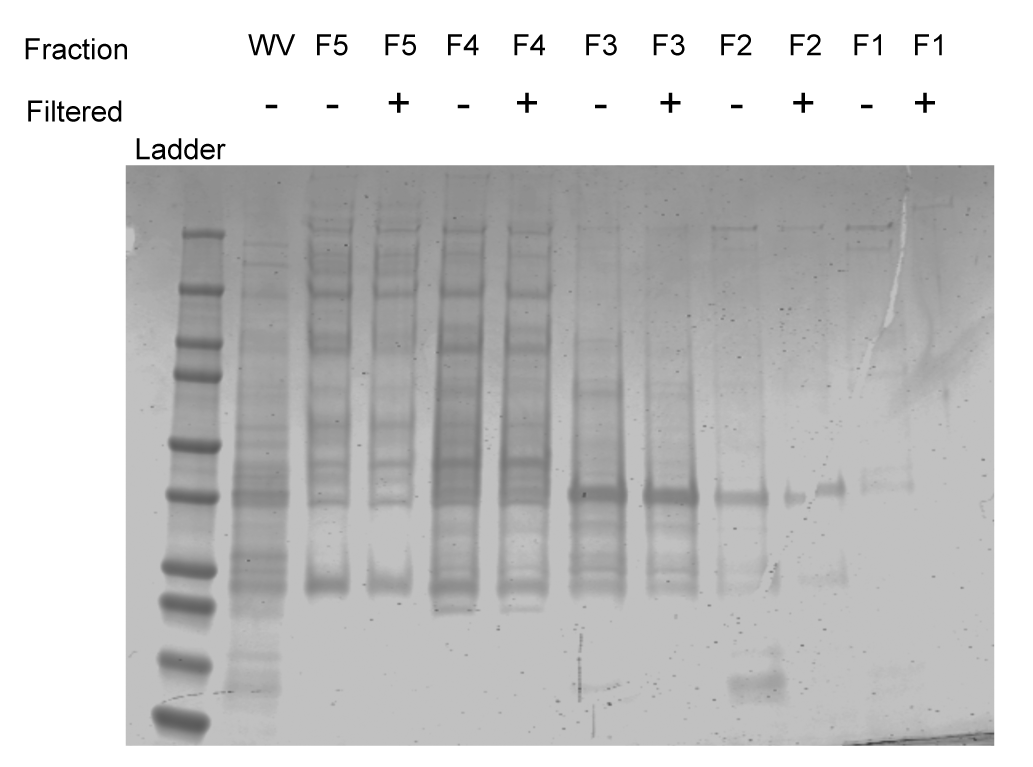
