## Supporting Information for "Venom vesicles from the parasitoid *Ganaspis hookeri* facilitate venom protein entry into host immune cells"

**Supporting Information Figure Legends**

**Figure S1**. Supporting information for Figure 1. Whole mount brightfield micrograph of the dissected venom apparatus shown in Figure 1A-C. Image is a single section taken at 40x magnification. Scale bar is 50 µm.

**Figure S2**. Supporting information for Figure 4. Additional venom vesicles from fraction F5. (A-B) Transmission electron micrographs of F5 vesicles displaying the unusual morphology. Scale bars are 100 nm in both panels.

**Figure S3**. Supporting information for Figure 7. Additional representative images of *hop[Tum]* cells incubated with Exo-Venom (A-B) and Exo-PBS (C-D). Confocal micrographs (A, C) shown with paired brightfield images (B, D).

**Figure S4**. Supporting information for Venom purification methods. SDS-PAGE analysis of fresh whole and sucrose gradient fractionated protein. Samples from each fraction were filtered (+) or not (-) prior to electrophoresis. Comparison of the banding patterns suggest that filtration does not remove any venom contents that are present in fresh venom. We can conclude that filtration allows us to remove aggregated material without selectively losing venom components.
